# Massively integrated coexpression analysis reveals transcriptional regulation, evolution and cellular implications of the noncanonical translatome

**DOI:** 10.1101/2023.03.16.533058

**Authors:** April Rich, Omer Acar, Anne-Ruxandra Carvunis

## Abstract

**Background:** Recent studies uncovered pervasive transcription and translation of thousands of noncanonical open reading frames (nORFs) outside of annotated genes. The contribution of nORFs to cellular phenotypes is difficult to infer using conventional approaches because nORFs tend to be short, of recent *de novo* origins, and lowly expressed. Here we develop a dedicated coexpression analysis framework that accounts for low expression to investigate the transcriptional regulation, evolution, and potential cellular roles of nORFs in *Saccharomyces cerevisiae*.

**Results:** Our results reveal that nORFs tend to be preferentially coexpressed with genes involved in cellular transport or homeostasis but rarely with genes involved in RNA processing. Mechanistically, we discover that young *de novo* nORFs located downstream of conserved genes tend to leverage their neighbors’ promoters through transcription readthrough, resulting in high coexpression and high expression levels. Transcriptional piggybacking also influences the coexpression profiles of young *de novo* nORFs located upstream of genes, but to a lesser extent and without detectable impact on expression levels. Transcriptional piggybacking influences, but does not determine, the transcription profiles of *de novo* nORFs emerging nearby genes. About 40% of nORFs are not strongly coexpressed with any gene but are transcriptionally regulated nonetheless and tend to form entirely new transcription modules. We offer a web browser interface (https://carvunislab.csb.pitt.edu/shiny/coexpression/) to efficiently query, visualize and download our coexpression inferences.

**Conclusions:** Our results suggest that nORF transcription is highly regulated. Our coexpression dataset serves as an unprecedented resource for unraveling how nORFs integrate into cellular networks, contribute to cellular phenotypes, and evolve.

## Background

Eukaryotic genomes encompass thousands of open reading frames (ORFs). The vast majority are so-called “noncanonical” ORFs (nORFs) excluded from genome annotations because of their short length, lack of evolutionary conservation, and perceived irrelevance to cellular physiology [1–3]. The development of RNA sequencing (RNA-seq) [4] and ribosome profiling [5,6] has revealed genome-wide transcription and translation of nORFs across species ranging from yeast to humans [6–14]. Recent studies have characterized individual nORFs that form stable peptides and impact phenotypes, including cell growth [10,13,15], cell cycle regulation [16], muscle physiology [17–19], and immunity [20–22]. Unraveling the cellular, physiological and evolutionary implications of nORFs has become an active area of research [14,23].

Many nORFs have evolved *de novo* from previously noncoding regions [24–26]. Thus, the study of nORFs and *de novo* gene birth as evolutionary innovation carries a synergistic overlap where findings in one area could improve our understanding of the other. For instance, Sandmann et al. measured physical protein interactions for hundreds of peptides translated from nORFs and proposed that short linear motifs present in young *de novo* nORFs could mediate how nORFs impact essential cellular processes [26]. Other studies observed a gradual integration of evolutionary young ORFs into cellular networks and showed they could gain essential roles [27–29]. These studies support an evolutionary model whereby pervasive expression of nORFs generates the raw material for *de novo* gene birth [24,25].

The biological interpretation of nORF expression is complex. Some studies suggest that the transcription or translation of nORFs could be attributed to expression noise [30–32], whereby non-specific binding of RNA polymerases and ribosomes to DNA and RNA might cause promiscuous transcription or translation, respectively. How do nORFs become expressed in the first place? There are multiple hypotheses on how *de novo* ORFs gain the ability to become transcriptionally regulated [33]. One possibility is the emergence of novel regulatory regions along with or following the emergence of an ORF (ORF-first), as was shown for specific *de novo* ORFs in *Drosophila melanogaster* [34], codfish [35], human [36,37] and chimpanzee [36]. Alternatively, ORFs may emerge on actively transcribed loci such as near enhancers [38] or on long noncoding RNAs [39], as was shown for *de novo* ORFs in primates [40] and for *de novo* ORFs upstream or downstream of transcripts containing genes [37] (transcription-first) [41–43]. Transcription has a ripple effect causing coordinated activation of nearby genes [44,45]. Thus, *de novo* ORFs that emerge near established genes or regulatory regions may acquire transcriptional regulation by ‘piggybacking’ [45] on the pre-existing regulatory context [41,46]. This piggybacking could predispose *de novo* ORFs to be involved in similar cellular processes as their neighbors, which in turn would help with characterization. To date, the fraction of nORFs that are transcriptionally regulated and contribute to cellular phenotypes is unknown for any species.

An obstacle to studying nORF expression at scale is their detection, as nORF expression levels are typically low and reliant on specific conditions [24,36]. Recent studies demonstrated that the integration of omics data [14,47–49] could effectively address detection issues. For example, Wacholder et al. [14] recently discovered around 19,000 translated nORFs in *Saccharomyces cerevisiae* by massive integration of ribosome profiling data. This figure is three times larger than the number of canonical ORFs (cORFs) annotated in the yeast genome. These translated nORFs have the potential to generate peptides that affect cellular phenotypes but are almost entirely uncharacterized.

Coexpression is a well-established approach for studying transcriptional regulation through the massive integration of RNA-seq data. Coexpression refers to the similarity between transcriptional profiles of ORF pairs across numerous samples. Coexpression has been used successfully to identify new gene functions [50,51], disease-related genes [22,52,53] and for studying the conservation of the regulatory machinery [51,54] or gene modules [55] between species. Based on the assumption that genes involved in similar pathways have correlated expression patterns, coexpression can reveal relationships between genes and other transcribed genetic elements [56,57]. Most coexpression studies have focused on cORFs, but the abundance of publicly available RNA-seq data represents a tractable avenue to interrogate the transcriptional regulation of thousands of nORFs at once using coexpression approaches [47,58–61]. Indeed, RNA-seq is probe-agnostic and annotation-agnostic, thereby enabling the reuse of existing data to explore these novel ORFs. However, low expression levels can distort coexpression inferences due to statistical biases [62,63]. A coexpression analysis of translated nORFs that addresses the statistical issues arising from low expression is still lacking for any species.

Here, we developed a dedicated statistical approach that accounts for low expression levels when inferring coexpression relationships between ORFs. We applied this approach to the recently identified 19,000 translated nORFs in *S. cerevisiae* [14] and built the first high-quality coexpression network spanning the canonical and noncanonical translatome of any species. Coexpression relationships suggest that the majority of nORFs are transcriptionally regulated. While many nORFs form entirely new noncanonical transcription modules, approximately half are transcriptionally associated with genes involved in cellular homeostasis and transport. We show that *de novo* ORFs that piggyback onto their neighbors’ transcription tend to have higher expression and tend to be highly coexpressed with their neighbors. We provide a web application to allow researchers to easily access this dataset to investigate the coexpression relationships and potential cellular roles for thousands of ORFs.

## Results

### High-quality coexpression inferences show transcriptional and regulatory relationships between nORFs and cORFs

To infer coexpression at the translatome scale in *S. cerevisiae*, we considered all cORFs annotated as “verified”, “uncharacterized”, or “transposable element” in the *Saccharomyces* Genome Database (SGD) [67], as well as all nORFs, ORFs that were either unannotated or annotated as “dubious” and “pseudogene”, with evidence of translation according to Wacholder et al. [14]. To maximize detection of transcripts containing nORFs, we curated and integrated 3,916 publicly available RNA-seq samples from 174 studies (Figure 1A, Supplementary Data 1). Many nORFs were not detected in most of the samples we collected, creating a very sparse dataset (Figure 1B). The issue of sparsity has been widely studied in the context of single cell RNA-seq (scRNA-seq). A recent study looking at multiple measures of association for constructing coexpression networks from scRNA-seq showed that proportionality methods coupled with center log ratio (clr) transformation consistently outperformed other measures of coexpression in a variety of tasks including identification of disease-related genes and protein-protein network overlap analysis [68]. Thus, we used clr to transform the raw read counts and quantified coexpression relationships using the proportionality metric, ρ [69].

**Figure 1:**
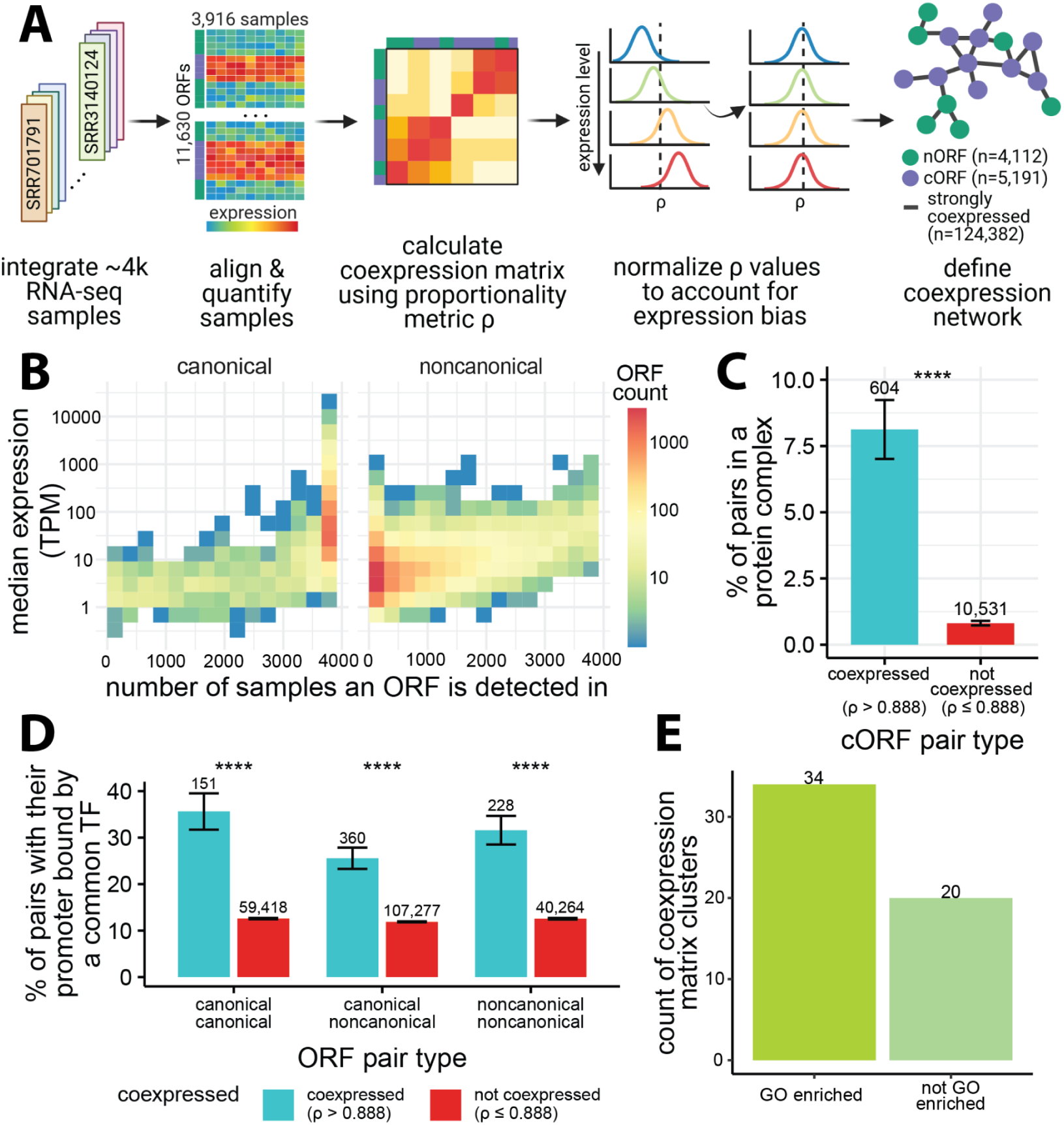
Overview of coexpression inference framework and properties of the dataset. A) Workflow: 3,916 samples were analyzed to create an expression matrix for 11,630 ORFs, including 5,803 cORFs and 5,827 nORFs; center log ratio transformed (clr) expression values were used to calculate the coexpression matrix using proportionality metric, ρ, followed by normalization to correct for expression bias. The coexpression matrix was thresholded using ρ > 0.888 to create a coexpression network (top 0.2% of all pairs). B) Distribution of the number of ORFs binned based on their median expression values (transcript per million - TPM) and the number of samples the ORFs were detected in with at least 5 raw counts. C) Coexpressed cORF pairs (ρ > 0.888) are more likely to encode proteins that form complexes than non-coexpressed cORF pairs (Fisher’s exact test p < 2.2e-16; error bars: standard error of the proportion); using annotated protein complexes from ref. [64]. D) Coexpressed ORF pairs (ρ > 0.888) are more likely to have their promoters bound by a common transcription factor (TF) than non-coexpressed ORF pairs (Fisher’s exact test p < 2.2e-16; error bars: standard error of the proportion); genome-wide TF binding profiles from ref. [65] and transcription start sites (TSS) from ref. [66] were analyzed to define promoter binding (see Methods). E) Hierarchical clustering of the coexpression matrix reveals functional enrichments for most clusters that contain at least 5 cORFs; functional enrichments estimated by gene ontology (GO) enrichment analysis at false discovery rate (FDR) < 0.05 using Fisher’s exact test.

We further addressed the issue of sparsity with two sample thresholding approaches. First, any observation with a raw count below five was discarded, such that when calculating ρ only the samples expressing both ORFs with at least five counts were considered. Second, we empirically determined that a minimum of 400 samples were required to obtain reliable coexpression values by assessing the effect of sample counts on the stability of ρ values (Supplementary Figure 1). These steps resulted in an 11,630 by 11,630 coexpression matrix encompassing 5,803 cORFs and 5,827 nORFs (ORF list in Supplementary Data 2).

The combined use of clr, ρ, and sample thresholding accounted for statistical issues in estimating coexpression deriving from sparsity, but the large difference in RNA expression levels between cORFs and nORFs posed yet another challenge. Indeed, Wang et al. showed that the distribution of coexpression values is biased by expression level due to statistical artifacts [62]. We observed this artifactual bias in our dataset (Supplementary Figure 2A) and corrected for it using spatial quantile normalization (SpQN) as recommended by Wang et al. [62] (Supplementary Figure 2B). This resulted in a normalized coexpression matrix (Supplementary Data 3) with ρ values centered around 0.476.

We then created a network representation of the coexpression matrix by considering only the top 0.2% of ρ values between all ORF pairs (ρ > 0.888). This threshold was chosen to include 90% of cORFs (Supplementary Figure 3). Altogether, our dedicated analysis framework (Figure 1A) inferred 124,382 strong (ρ > 0.888) coexpression relationships between 9,303 ORFs, encompassing 4,112 nORFs and 5,191 cORFs.

To assess whether our coexpression network captures meaningful biological and regulatory relationships, we examined its overlap with orthogonal datasets. Using a curated [64] protein complex dataset for cORFs, we found that coexpressed cORF pairs are significantly more likely to encode proteins that form a protein complex together compared to non-coexpressed pairs (Odds ratio = 10.8 Fisher’s exact test p < 2.2e-16; Figure 1C). Using a previously published [65] genome-wide chromatin immunoprecipitation with exonuclease digestion (ChIP-exo) dataset containing DNA-binding information for 73 sequence-specific transcription factors (TFs) and using transcript isoform sequencing (TIF-seq) [66] data to determine transcription start sites (TSSs) and promoter regions, we observed that coexpressed ORF pairs were more likely to have their promoters bound by a common TF than non-coexpressed ORF pairs, whether the pairs consist of nORFs or cORFs (*canonical-canonical pairs*: Odds ratio = 3.84, *canonical-noncanonical pairs*: Odds ratio = 2.55, *noncanonical-noncanonical pairs*: Odds ratio = 3.22, Fisher’s exact test p < 2.2e-16 for all three comparisons; Figure 1D). Enrichments were robust to different coexpression cutoffs (Supplementary Figure 4-5). Using the WGCNA [70] method to cluster the coexpression matrix, we found that more than half of the clusters identified contained functionally related ORFs (gene ontology (GO) biological process enrichments at Benjamini-Hochberg (BH) adjusted false discovery rate (FDR) < 0.05; Figure 1E; Supplementary Figure 6). These analyses demonstrate the high quality of our coexpression network and confirm that it captures meaningful biological and regulatory relationships for both cORFs and nORFs.

Conventional approaches for coexpression analysis include using transcript per million (TPM) or reads per kilobase per million (RPKM) normalization, batch correction by removing top principal components, and Pearson’s correlation as the similarity metric [71,56,72]. Compared to these approaches, our framework increased the proportion of coexpressed ORF pairs whose promoters are bound by a common TF specifically for pairs containing nORFs (Supplementary Figure 7), and yielded coexpression networks encompassing the largest number of nORFs at most thresholds (Supplementary Figure 8). Hence our dedicated analysis framework therefore outperforms conventional coexpression approaches for the study of nORFs. We offer an R Shiny [73] interface (https://carvunislab.csb.pitt.edu/shiny/coexpression/) to efficiently query, visualize and download the coexpression data we generated. To our knowledge, this is the most comprehensive coexpression dataset focusing on empirically translated elements, both annotated and unannotated, for any species to date.

### nORFs tend to be located at the periphery of the coexpression network and form new noncanonical transcription modules

Conventional analyses of coexpression networks have been restricted to cORFs. Our full coexpression network contains twice the number of ORFs and three times the number of strong (ρ > 0.888) coexpression relationships compared to the canonical-only network (Figure 2A). We sought to compare the network properties of the canonical-only and full networks. On average, nORFs have fewer coexpressed partners (degree) than cORFs, suggesting that nORFs have distinct transcriptional profiles (Cliff’s Delta d = −0.29, Mann-Whitney U-test p < 2.2e-16; Figure 2B). We found that 91% of cORFs are coexpressed with at least one nORF (n = 4,726; Figure 2C), whereas only 59% of nORFs are coexpressed with at least one cORF. In contrast, we would have expected an average of 89% of nORFs to be coexpressed with a cORF according to degree preserving simulations of 1,000 randomized networks where edges from nORFs were shuffled (Odds ratio = 0.174, Fisher’s exact test p < 2.2e-16; Figure 2D, Supplementary Figure 9). This suggests that, while most nORFs are integrated in the full coexpression network, they also have distinct expression profiles that differ markedly from those of all cORFs and are more similar to those of other nORFs.

**Figure 2.**
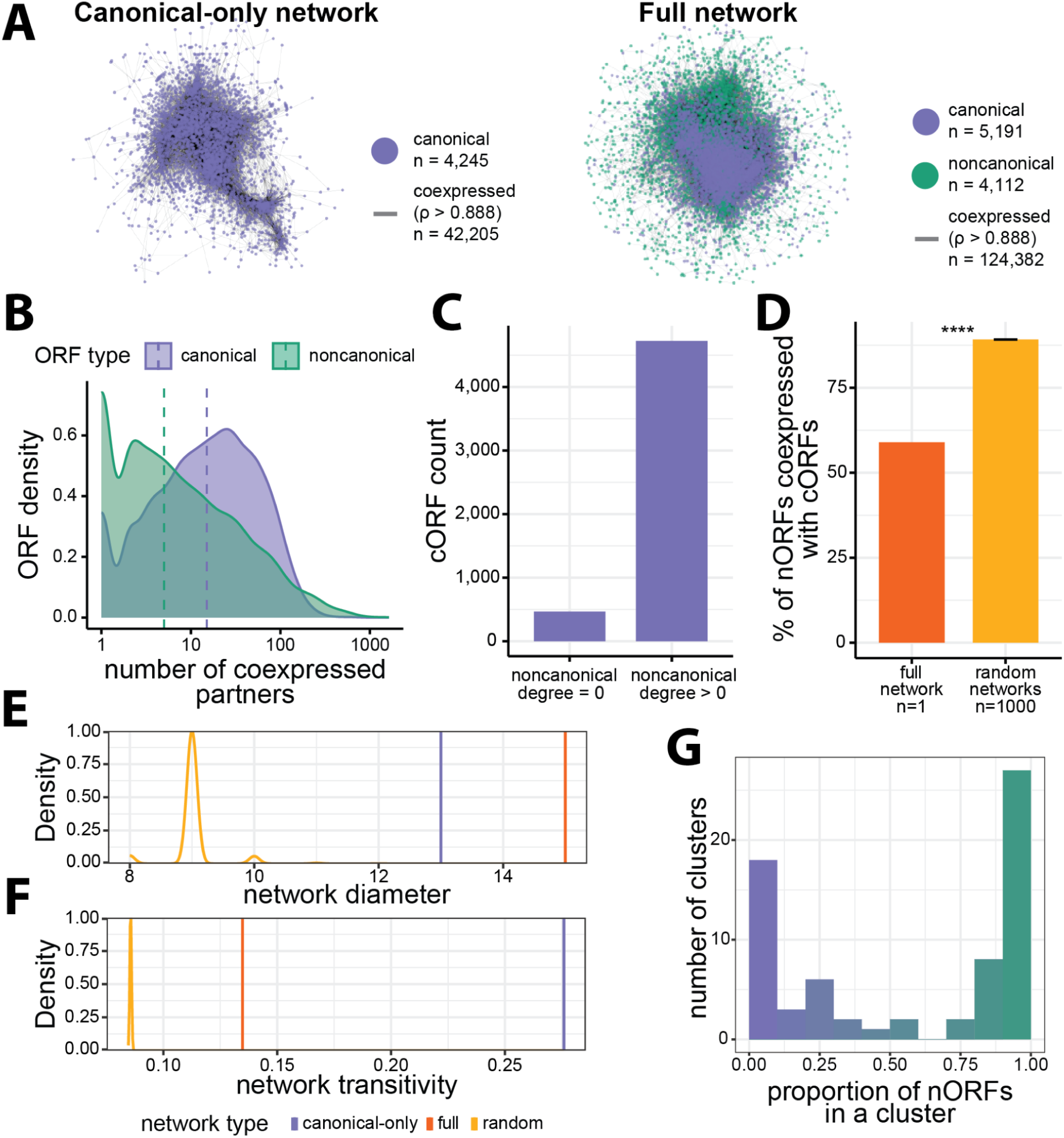
Topological properties of the coexpression network. A) Visualization for canonical-only and full coexpression networks using spring embedded graph layout [74]. The full network contains more cORFs than the canonical-only network since addition of nORFs also results in addition of many cORFs that are only connected to an nORF. B) nORFs have fewer coexpression partners (degree in full network) than cORFs (Mann-Whitney U-test p < 2.2e-16). C) Most cORFs are coexpressed with at least one nORF. D) Only 59% of nORFs are coexpressed with at least one cORFs and this is less than expected by chance, on average, 89% of nORFs are coexpressed with a cORF across 1,000 randomized networks generated in a degree-preserving fashion by swapping edges of noncanonical nodes (Fisher’s exact test p < 2.2e-16; error bar: standard error of the mean proportion across randomized networks). E) Addition of nORFs to the canonical-only network results in the full network being less compact, whereas the opposite is expected by chance, shown by the decrease in diameters for the 1,000 randomized networks. F) Addition of nORFs to the canonical-only network decreases local clustering in the full network, however this is to a lesser extent than expected by chance as shown by the distribution for the 1,000 randomized networks. G) Most clusters in the coexpression matrix encompass either primarily nORFs or primarily cORFs (n= 69 clusters, *green* represents nORF majority clusters, *purple* represents cORF majority clusters).

To investigate how these seemingly conflicting attributes impact the organization of the coexpression network, we analyzed two global network properties: diameter, which is the longest shortest path between any two ORFs; and transitivity, which is the tendency for ORFs that are coexpressed with a common neighbor to also be coexpressed with each other. The incorporation of nORFs in the full network led to a larger diameter relative to the canonical-only network (Figure 2E). This is in sharp contrast with the null expectation, set by 1,000 degree-preserving simulations, whereby random incorporation of nORFs decreases network diameter. The full coexpression network is thus much less compact than expected by chance, suggesting that nORFs tend to be located at the periphery of the network. Network transitivity decreased with the incorporation of nORFs compared to the canonical-only network, but to a lesser extent than expected by chance (Figure 2F). This suggests that despite their low degree and peripheral locations, the connections formed by nORFs are structured and may form noncanonical clusters.

To investigate this hypothesis, we inspected the ratio of nORFs and cORFs among the cluster assignments from WGCNA hierarchical clustering of the full coexpression matrix (Supplementary Figure 6). Strikingly, we observed a bimodal distribution of clusters, with approximately half of the clusters consisting mostly of nORFs and the other half containing mostly cORFs (Figure 2G). We conclude that nORFs exhibit a unique and non-random organization within the coexpression network, simultaneously connecting to all cORFs while also forming entirely new noncanonical transcription modules.

### Coexpression profiles reveal most nORFs are transcriptionally associated with genes involved in cellular transport and homeostasis

To determine whether nORFs are transcriptionally associated with specific cellular processes, we performed gene set enrichment analyses [77] (GSEA) on their coexpression partners. GSEA takes an ordered list of genes, in this case sorted by coexpression level, and seeks to find if the higher ranked genes are preferentially annotated with specific GO terms. For each cORF and nORF, we ran GSEA to detect if their highly coexpressed partners were preferentially associated with any GO terms (Supplementary Figure 10). Almost all ORFs (99.9%), whether cORF or nORF, had at least one significant GO term associated with their coexpression partners at BH adjusted FDR < 0.01, suggesting that nORFs are engaged in coherent transcriptional programs. We then calculated, for each GO term, the number of cORFs and nORFs with GSEA enrichments in this term (Supplementary Data 4). These analyses identified specific GO terms that were significantly more (16 terms, BH adjusted FDR < 0.001, Odds ratio > 2, Fisher’s exact test; Figure 3A, Supplementary Data 5) or less (23 terms, BH adjusted FDR < 0.001, Odds ratio < 2, Fisher’s exact test; Figure 3B, Supplementary Data 5) prevalent among the coexpression partners of nORFs relative to those of cORFs. Most of the GO terms that were significantly enriched among the coexpression partners of nORFs were related to cellular homeostasis and transport (Figure 3A) while most of the GO terms significantly depleted among the coexpression partners of nORFs were related to DNA, RNA, and protein processing (Figure 3B). Running the same GSEA pipeline with Kyoto Encyclopedia of Genes and Genomes (KEGG) [78] annotations yielded consistent results (Supplementary Figure 11, Supplementary Data 6-7). Half of nORFs were coexpressed with genes involved in homeostasis (GO:0042592, 53%), monoatomic ion transport (GO:0006811, 49%) and transmembrane transport (GO:0055085, 47%). The nORFs transcriptionally associated with the parent term ‘transport’ (n = 2,718, GO:0006810, GSEA BH adjusted FDR < 0.01) were 1.6 times more likely to contain a predicted transmembrane domain than other nORFs (p = 1.3e-4, Fisher’s exact test; Figure 3C), in line with potential transport-related activities. These findings reveal a strong and previously unsuspected transcriptional association between nORFs, and cellular processes related to homeostasis and transport.

**Figure 3.**
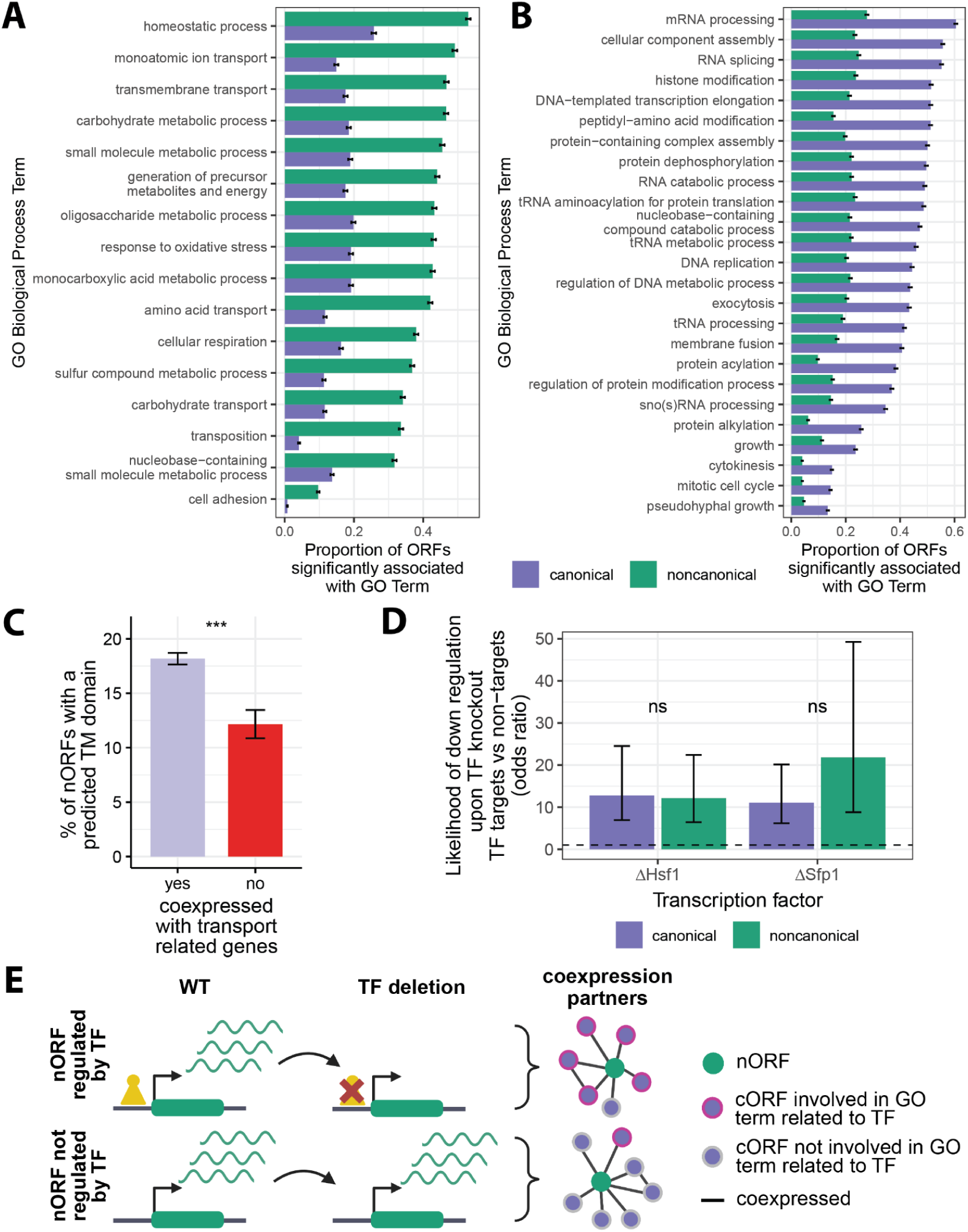
Biological processes associated with nORF transcriptional regulation. A-B) Biological processes that are more (A) (Odds ratio > 2, n = 16 terms) or less (B) (Odds ratio < 0.5, n = 23 terms) transcriptionally associated with nORFs than cORFs (y-axis ordered by nORF enrichment proportion from highest to lowest, BH adjusted FDR < 0.001 for all terms, Fisher’s exact test, GO term enrichments were detected using gene set enrichment analyses (GSEA), error bars: standard error of the proportion). C) nORFs that are highly coexpressed with genes involved in transport are more likely to have predicted transmembrane (TM) domains as determined by TMHMM [75] compared to nORFs that are not (Odds ratio = 1.6, Fisher’s exact test p = 1.3e-4; error bars: standard error of the proportion). D) nORFs and cORFs that are Sfp1 or Hsf1 targets are more likely to be downregulated when Sfp1 or Hsf1 are deleted compared to ORFs that are not targets (*Sfp1*: cORFs: p < 2.2e-16; nORFs: p = 2.8e-9; *Hsf1*: cORFs: p <2.2e-16; nORFs: p = 9.9e-13; Fisher’s exact test, error bars: 95% confidence interval of the odds ratio; *dashed* line shows odds ratio of 1; RNA abundance data from SRA accession SRP159150 and SRP437124 [76] respectively). E) nORFs that are regulated by TFs are more likely to be coexpressed with genes involved in processes related to known functions of that TF.

### Hsf1 and Sfp1 nORF targets are part of protein folding and ribosome biogenesis transcriptional programs, respectively

Overall, our analyses relating coexpression to TF binding (Figure 1D) and functional enrichments (Figure 3A-B) suggest that nORF expression is regulated rather than simply the consequence of transcriptional noise. To further investigate this hypothesis, we sought to identify regulatory relationships between specific TFs and nORFs. We reasoned that if nORFs are regulated by TFs in similar ways as cORFs, then genetic knockout of the TFs that regulate them should impact their expression levels as it does for cORFs [79]. We focused on two transcriptional activators for which both ChIP-exo [65] and knockout RNA-seq data [76] were publicly available: Sfp1, which regulates ribosome biogenesis [80] and Hsf1, which regulates heat shock and protein folding responses [81].

For both cORFs and nORFs, knockout of Sfp1 or Hsf1 was more likely to trigger a significant decrease in expression when the ORF’s promoter was bound by the respective TF according to ChIP-exo evidence (Figure 3D). The statistical association between TF binding and knockout-induced downregulation was as strong for nORFs as it was for cORFs, consistent with nORFs having similar mechanisms of transcriptional activation (*Sfp1*: cORFs Odds ratio = 11.1, p < 2.2e-16; nORFs Odds ratio = 21.8, p = 2.8e-9, Fisher’s exact test; *Hsf1*: cORFs Odds ratio = 12.7, p < 2.2e-16; nORFs Odds ratio = 12.1, p = 9.9e-13, Fisher’s exact test). Therefore, the nORFs whose promoters are bound by these TFs, and whose expression levels decrease upon deletion of these TFs, are likely genuine regulatory targets of these TFs. By this stringent definition, our analyses identified 9 nORF targets of Sfp1 (and 34 cORF targets) and 19 nORF targets of Hsf1 (and 39 cORF targets). The coexpression profiles of these Sfp1 and Hsf1 nORF targets were preferentially associated with genes involved in processes directly related to the known functions of Sfp1 and Hsf1 (Supplementary Data 8). For example, the coexpression profiles of 9 Sfp1 nORF targets revealed preferential associations with genes involved in ‘ribosomal large subunit biogenesis’ and 7 Sfp1 nORF targets involved in ‘regulation of translation’ according to our GSEA pipeline (Fisher’s exact test, BH adjusted p-value < 6.7e-4 for both terms). Similarly, 13 Hsf1 nORF targets were preferentially associated with genes involved in ‘Protein Folding’ (Fisher’s exact test, BH adjusted p-value = 5.7e-9). These results show that nORF expression can be actively regulated by TFs as part of coherent transcriptional programs (Figure 3E).

### *de novo* ORF expression and regulation are shaped by genomic location

Previous literature has shown that many nORFs arise *de novo* from previously noncoding regions [24,26]. We wanted to investigate how these evolutionarily novel ORFs acquire expression and whether their locus of emergence influences this acquisition. To define which ORFs were of recent *de novo* evolutionary origins, we developed a multistep pipeline combining sequence similarity searches and syntenic alignments (Figure 4A). cORFs were considered conserved if they had homologues detectable by sequence similarity searches with BLAST in budding yeasts outside of the *Saccharomyces* genus or if their open reading frames were maintained within the *Saccharomyce*s genus [14]. cORFs and nORFs were considered *de novo* if they lacked homologues detectable by sequence similarity outside of the *Saccharomyces* genus and if less than 60% of syntenic orthologous nucleotides in the two most distant *Saccharomyces* branches were in the same reading frame as in *S. cerevisiae*. These criteria aimed to identify the youngest *de novo* ORFs. Overall, we identified 5,624 conserved cORFs and 2,756 *de novo* ORFs including 77 *de novo* cORFs and 2,679 *de novo* nORFs (Figure 4B). In general, the coexpression patterns of *de novo* ORFs (Supplementary Figure 12) were similar to those of nORFs (Figure 3A-B).

**Figure 4.**
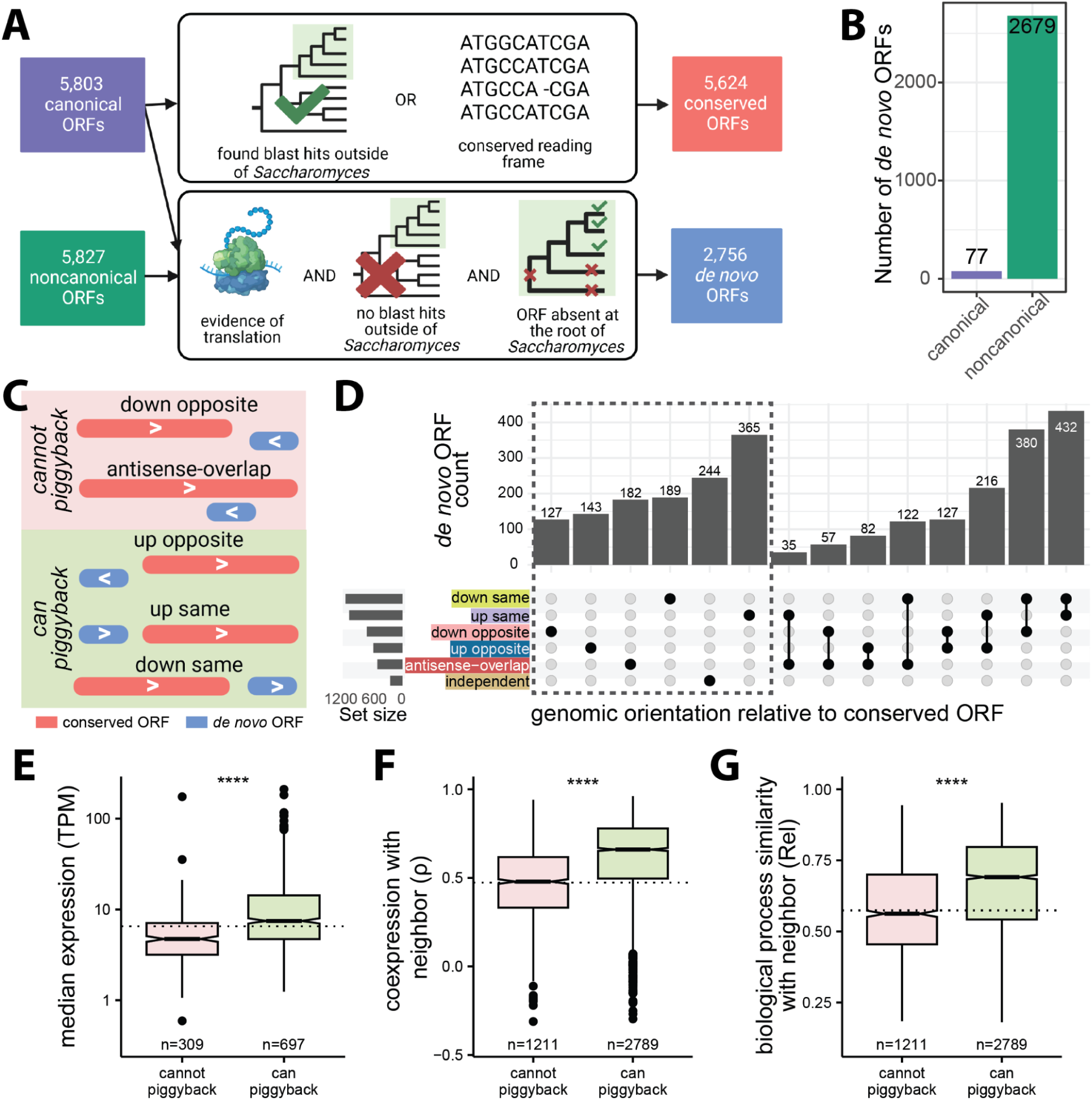
Expression, coexpression and biological processes similarity of *de novo* ORFs with respect to genomic orientations. A) Pipeline used to reclassify ORFs as conserved or *de novo*. cORFs were considered for both conserved and *de novo* classification while nORFs were only considered for *de novo* classification. Conserved ORFs were determined by either detection of homology outside of *Saccharomyces* or reading frame conservation within *Saccharomyces* (*top*). *De novo* ORFs were determined by evidence of translation, lack of homology outside of *Saccharomyces* as well as lack of a homologous ORF in the two most distant *Saccharomyces* branches (*bottom*). B) Counts of cORFs and nORFs that emerged *de novo*. C) Genomic orientations of *de novo* ORFs that cannot transcriptionally piggyback off neighboring conserved ORF (cannot share promoter with neighbor, *pink shading*) or can transcriptionally piggyback off neighboring conserved ORF (possible to share promoter with neighbor, *green shading*). D) Counts of *de novo* ORFs that are within 500 bp of a conserved ORF in different genomic orientations; ORFs further than 500bp are classified as independent. E) *De novo* ORFs in orientations that can piggyback have higher RNA expression levels than *de novo* ORFs in orientations that cannot piggyback (Cliff’s Delta d = 0.4). Only *de novo* ORFs in a single orientation are considered (dashed box in panel *D*). Dashed line represents the median expression of independent *de novo* ORFs. F) *de novo* ORFs in orientations that can piggyback have higher coexpression with neighboring conserved ORFs compared to *de novo* ORFs in orientations that cannot piggyback (Cliff’s Delta d = 0.43). Dashed line represents median coexpression of *de novo*-conserved ORF pairs on separate chromosomes. G) *de novo* ORFs in orientations that can piggyback are more likely to be transcriptionally associated with genes involved in the same biological processes as their neighboring conserved ORFs than *de novo* ORFs in orientations that cannot piggyback (Cliff’s Delta d = 0.31). Dashed line represents median functional enrichment similarities of *de novo*-conserved ORF pairs on separate chromosomes. (For panels E-F-G: Mann-Whitney U-test, ****: p < 2.2e-16).

We hypothesized that the locus where *de novo* ORFs arise may influence their expression profiles through “piggybacking” off their neighboring conserved ORFs’ pre-existing regulatory environment. To investigate this hypothesis, we categorized *de novo* ORFs based on their positioning relative to neighboring conserved ORFs. The *de novo* ORFs further than 500 bp from all conserved ORFs were classified as independent. The remaining *de novo* ORFs were classified as either upstream or downstream on the same strand (up same or down same), upstream or downstream on the opposite strand (up opposite or down opposite), or as overlapping on the opposite strand (anti-sense overlap) based on their orientation to the nearest conserved ORF (Figure 4C-D). We categorized the orientations as being able to piggyback or unable to piggyback based on their potential of sharing a promoter with neighboring conserved ORFs, with down opposite and antisense overlap as orientations that cannot piggyback and up opposite, up same, and down same as orientations that can piggyback (Figure 4C). The piggybacking hypothesis predicts that *de novo* ORFs that arise in orientations that can piggyback would be positively influenced by the regulatory environment provided by the promoters of neighboring conserved ORFs, resulting in similar transcription profiles as their neighbors and increased expression relative to *de novo* ORFs that do not benefit from a pre-existing regulatory environment.

We considered three metrics to assess piggybacking: RNA expression level, measured as median TPM over all the samples analyzed, coexpression with neighboring conserved ORF and biological process similarity with neighboring conserved ORF. To calculate biological process similarity between two ORFs, we used significant GO terms at FDR < 0.01 determined by coexpression GSEA for each ORF (Supplementary Figure 10) and calculated the similarity between these two sets of GO terms using the relevance method [82]. If two ORFs are enriched in the same specialized terms, their relevance metric would be higher than if they are enriched in different terms or in the same generic terms. We found that *de novo* ORFs in orientations that can piggyback tend to have higher expression (focusing only on ORFs that could be assigned a single orientation, dashed box in Figure 4D, Cliff’s Delta d = 0.4; Figure 4E), higher coexpression with their neighbor (Cliff’s Delta d = 0.43; Figure 4F), and higher biological process similarity (Cliff’s Delta d = 0.31; Figure 4G), compared to ORFs in orientations that cannot piggyback (p < 2.2e-16 Mann-Whitney U-test for all). Thus, all three metrics supported the piggybacking hypothesis.

Closer examination revealed a more complex situation. First, the immediate neighbors of *de novo* ORFs in orientations that can piggyback were rarely among their strongest coexpression partners (only found in the top 10 coexpressed partners for 15% of down same, 4.5% of up same, 3% of up opposite ORFs). Therefore, emergence nearby a conserved ORF in a piggybacking orientation influences, but does not fully determine, the transcription profiles of *de novo* ORFs. Transcriptional regulation beyond that provided by the pre-existing regulatory environment may exist. Second, while ORFs in all three orientations that can piggyback displayed increased coexpression and biological process similarity with their neighbors relative to background expectations (Supplementary Figure 13A-B), only down same *de novo* ORFs displayed increased RNA expression levels (Supplementary Figure 13C). The expression levels of up same *de novo* ORFs were statistically indistinguishable from independent *de novo* ORFs, while those of up opposite *de novo* ORFs were significantly lower than those of independent *de novo* ORFs (Supplementary Figure 13C). Down same *de novo* ORFs also showed stronger coexpression and biological process similarity with their conserved neighbors than up same and up opposite *de novo* ORFs (Supplementary Figure 13A-B). Therefore, the transcription of down same *de novo* ORFs appeared most susceptible to piggybacking.

To understand the molecular mechanisms leading to the differences in expression, coexpression and biological process similarity between the orientations that can piggyback, which all have the potential to share a promoter with neighboring conserved ORF, we investigated which actually do by analyzing transcript architecture. Using a publicly available TIF-seq dataset [66], we defined down same or up same ORFs as sharing a promoter with their neighbor if they mapped to the same transcript at least once. We defined up opposite ORFs as sharing a promoter with their neighbor if their respective transcripts did not have overlapping TSSs, as would be expected for divergent promoters [83]. According to these criteria, 84% of down same (n = 174), 64% of up same (n = 368), and 66% of up opposite (n = 185) *de novo* ORFs share a promoter with their neighboring conserved ORFs (Supplementary Figure 14). Among all *de novo* ORFs that arose in orientations that can piggyback, those that share promoters with neighboring conserved ORFs displayed higher expression levels than those that do not (*down same*: d = 0.75, p = 1.06e-8; *up same*: d = 0.38, p = 1.23e-7; *up opposite*: d = 0.3, p = 2.9e-3 Mann-Whitney U-test, d: Cliff’s Delta; Figure 5A). We also observed a significant increase in coexpression and biological process similarity between *de novo* ORFs and their neighboring conserved ORFs when their promoters are shared compared to when they are not (coexpression: *down same*: d = 0.28, p = 2.99e-9; *up same*: d = 0.31, p < 2.2e-16; *up opposite*: d = 0.27, p = 2.1e-7; biological process similarity: *down same*: d = 0.24, p = 5.5e-7; *up same*: d = 0.108, p = 3.78e-3; *up opposite*: d = 0.24, p = 6.1e-6, d: Cliff’s Delta, Mann-Whitney U-test; Figures 5B and 5C, respectively). Hence, sharing a promoter led to increases in the three piggybacking metrics for the three orientations.

**Figure 5.**
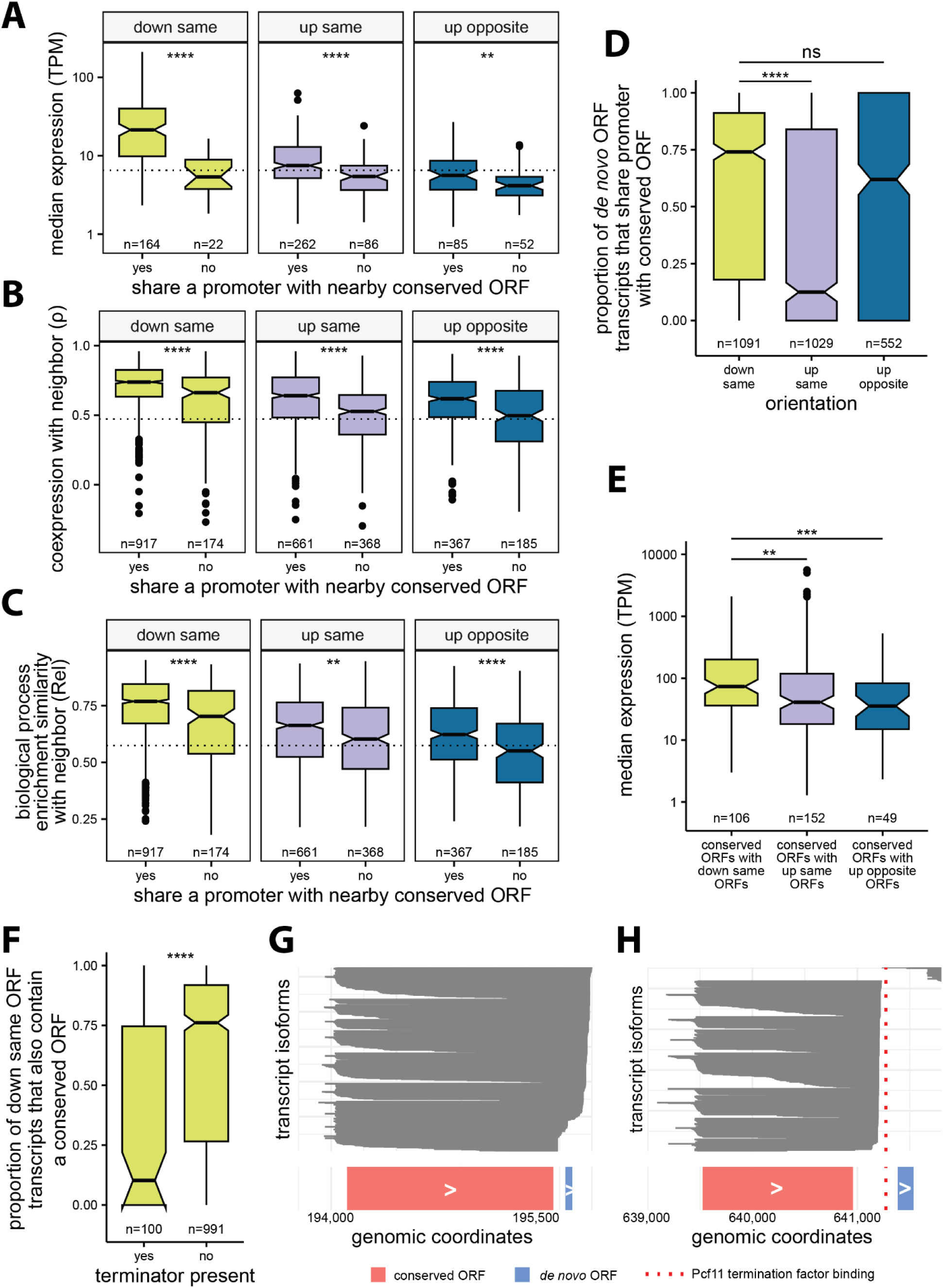
Effects of promoter sharing on expression, coexpression and biological process similarities of *de novo* ORFs. A) *De novo* ORFs that share a promoter with neighboring conserved ORFs, as determined by TIF-seq transcript boundaries, have significantly higher expression levels than *de novo* ORFs that do not. Considering only ORFs in a single orientation. Dashed line represents the median expression of independent *de novo* ORFs. B) *De novo* ORFs that share a promoter with neighboring conserved ORFs have higher coexpression with their neighbors than *de novo* ORFs that do not share a promoter. Dashed line represents median coexpression of *de novo*-conserved ORF pairs on separate chromosomes. C) *De novo* ORFs that share a promoter have more similar functional enrichments with neighboring conserved ORFs than *de novo* ORFs that do not share a promoter. Dashed line represents median functional enrichment similarities of the background distribution of *de novo*-conserved ORF pairs on separate chromosomes. D) Down same *de novo* ORFs share a promoter with neighboring conserved ORFs significantly more often than up same ORFs. E) Conserved ORFs with downstream *de novo* ORFs have a significant increase in expression compared to conserved ORFs with upstream *de novo* ORFs. F) Existence of transcription termination factors (Pcf11 or Nrd1) in between conserved ORFs and nearby downstream *de novo* ORFs leads to less shared transcripts. G) Transcript isoforms (*gray*) at an example locus where there are no transcription termination factors present between conserved ORF YBL015W (*pink*) and downstream *de novo* ORF chr2:195794-195847(+) (*blue*). H) Transcript isoforms (*gray*) at an example locus where there is Pcf11 transcription terminator present (*red line*) between conserved ORF YPR034W (*pink*) and downstream *de novo* ORF chr16:641385-641534(+) (*blue*). All detected transcript isoforms on these loci are plotted for G and F. (For all panels: ****: p ≤ 0.0001, ***: p ≤ 0.001, **: p ≤ 0.01, *: p ≤ 0.05, ns: not-significant; Mann-Whitney U-test)

Further supporting the notion that down same ORFs are particularly prone to piggybacking, the down same *de novo* ORFs that share a promoter with their conserved neighbors displayed much higher expression levels, and higher coexpression and biological process similarity with their conserved neighbor, than up same or up opposite ORFs that also share a promoter with their conserved neighbors (expression: *down same vs up same*: d = 0.58; *down same vs up opposite*: d = 0.55; coexpression: *down same vs up same:* d = 0.29, *down same vs up opposite:* d = 0.38; biological process similarity: *down same vs up same*: d = 0.37, *down same vs up opposite*: d = 0.45; d: Cliff’s Delta, p < 2.2e-16 for all comparisons, Mann-Whitney U-test). This could be due to down same ORF’s tendency to share promoters more often than up same ORFs, as a larger proportion of transcripts containing down same ORFs also contain a conserved ORF (*down same vs up same*: Cliff’s Delta d = 0.26, Mann-Whitney U-test p < 2.2e-16; Figure 5D), or higher expression levels of conserved ORFs that have down same ORFs on their transcripts compared to conserved ORFs with up same or up opposite piggybacking ORFs (*down same vs up same*: d = 0.2, p = 5.4e-3; *down same vs up opposite*: d = 0.34, p = 6.5e-4, Mann-Whitney U-test, d: Cliff’s Delta; Figure 5E).

Based on these results, we reasoned that transcriptional readthrough could be the molecular mechanism underlying the efficient transcriptional piggybacking of down same *de novo* ORFs. To investigate this hypothesis, we examined the impact of transcription terminators Pcf11 or Nrd1 on the frequency of transcript sharing between a conserved ORF and its downstream *de novo* ORF. Analyzing publicly available ChIP-exo data [65], we found that the presence of terminators between conserved ORFs and their downstream *de novo* ORF pairs resulted in a notably lower percentage of shared transcripts (Cliff’s Delta d = −0.39, p = 1.59e-10, Mann-Whitney U-test; Figure 5F). As an illustration, consider the genomic region on chromosome II from bases 194,000 to 196,000, containing the conserved ORF YBL015W and a downstream *de novo* ORF (positions 195,794 to 195,847). No terminator factor is bound to the intervening DNA between these two ORFs. This pair has high coexpression, with ρ = 0.96 and we observed that nearly all transcripts in this region containing the *de novo* ORF also include YBL015W (Figure 5G). In contrast, the genomic region on chromosome XVI from 639,000 to 641,800, containing the conserved ORF YPR034W and downstream *de novo* ORF (positions 641,385 to 641,534), does have a Pcf11 terminator factor between the pair, and as expected, none of the transcripts in this region contain both YPR034W and the *de novo* ORF, which have poor coexpression as a result (ρ = 0.1; Figure 5H). We conclude that sharing a transcript via transcriptional readthrough is the major transcriptional piggybacking mechanism for down same *de novo* ORFs.

## Discussion

We explored the transcription of nORFs from multiple angles including network topology, associations with cellular processes, TF regulation, and influence of the locus of emergence on *de novo* ORF expression. Delving into network topology, we find that nORFs have distinct expression profiles that are strongly correlated with only a few other ORFs. Nearly all cORFs are coexpressed with at least one nORF, but the converse is not true. Numerous nORFs form new structured transcriptional modules, possibly involved in both known and unknown cellular processes. The addition of nORFs to the cellular network resulted in a more clustered network than expected by chance, highlighting the previously unsuspected influence of nORFs in shaping the coexpression landscape.

Our study is the first to show a large-scale association between the expression of nORFs and cellular homeostasis and transport processes. We anticipate that future studies will follow up to test these associations experimentally. We also found nORFs to be preferentially associated with cellular processes related to metabolism, transposition and cell adhesion, but rarely with the core processes of the central dogma, DNA, RNA or protein processing. Genes involved in transport, metabolism, and stress tend to have more variable expression compared to genes in other pathways [84]. Pathways with more variable expression could be more likely to incorporate novel ORFs, possibly as a form of adaptive transcriptional response. There are several consistent observations in the literature [47,85,86]. For instance, Li et al. [47] showed that many *de novo* ORFs are upregulated in heat shock. Wilson and Masel [87] found higher translation of *de novo* ORFs under starvation conditions. Carvunis et al. [24] found *de novo* cORFs are enriched for the GO term ‘response to stress’. Other studies showed examples of how specific *de novo* ORFs could be involved in stress response [35,88] or homeostasis [88,89]. For instance the *de novo* antifreeze glycoprotein AFGP allows Arctic codfish to live in colder environments [35] or *MDF1* in yeast [88,90] was found in a screen to provide resistance to certain toxins and mediates ion homeostasis [91]. Our results, combined with these previous investigations, argue that a large fraction of nORFs provide adaptation to stresses and help maintain homeostasis, perhaps through modulation of transport processes.

Recent research in yeast has revealed an enrichment of transmembrane domains [15,24,92,93] within *de novo* ORFs. Previous studies identified small nORFs and *de novo* ORFs that localize to diverse cellular membranes, such as those of the endoplasmic reticulum, Golgi, or mitochondria in different species [10,15,94–97]. These findings are consistent with the notion that *de novo* ORFs could play a role in a range of transport processes, such as ion, amino acid, or protein transport across cellular membranes. By establishing a connection between predicted transmembrane domains and increased coexpression with transport-related genes, our findings set the stage for future experimental investigations into the precise molecular mechanisms and functional roles of nORFs in diverse transport systems.

Lastly, we explored how the preexisting regulatory context influences the transcriptional profiles of *de novo* ORFs. We found that *de novo* ORFs that piggyback off their neighboring conserved ORFs’ promoters had increases in expression, coexpression and biological process similarity with their neighboring conserved ORFs. Strikingly, ORFs that emerge *de novo* downstream of conserved ORFs have the largest increases in expression, coexpression and biological process similarities with their neighbors compared to other orientations, largely due to transcriptional readthrough leading to transcript sharing. Previous studies have shown that the transcription of regions downstream of genes is functional and regulated [98]. A study in humans showed that readthrough transcription downstream of some genes is responsible for roughly 15%–30% of intergenic transcription and is induced by osmotic and heat stress creating extended transcripts that play a role in maintaining nuclear stability during stress [99]. Another study in humans and zebrafish showed that the translation of small ORFs located in the 3’ UTR of mRNAs (dORFs) increased the translation rate of the upstream gene [100]. Lastly, a study in yeast found that genes which are preferentially expressed as bicistronic transcripts tend to contain evolutionarily younger genes compared to adjacent genes that do not share transcripts, suggesting that transcript sharing could provide a route for novel ORFs to become established genes [101]. These findings together with our results suggest that genomic regions downstream of genes may provide the most favorable environment for the transcription of *de novo* ORFs.

Our analyses show that the likelihood of a *de novo* ORF being expressed or repressed under the same conditions as the neighboring conserved ORF is influenced by the extent to which it piggybacks on the neighboring ORF’s regulatory context. Therefore, in addition to the evolutionary pressure acting on the sequence of emerging ORFs, our results suggest that transcriptional regulation and genomic context also influence their functional potential. However, this influence is not entirely deterministic, and much weaker when *de novo* ORFs emerge upstream than downstream of genes. Future studies are needed to map regulatory networks controlling nORF expression and reconstruct their evolutionary histories.

There are several limitations to our study. First, while SpQN enhances the coexpression signal of lowly expressed ORFs, it comes at the cost of reducing signals in highly expressed ORFs [62]. Given our objective of studying lowly-expressed nORFs this tradeoff is deemed worthwhile. Second, our study provides evidence of associations between nORFs and cellular processes such as homeostasis and transport, but these findings are based on transcription profile similarities which do not necessarily imply cotranslation or correlated protein abundances [102]. Furthermore, our analyses were performed in the yeast *S. cerevisiae* and the generalizability of our findings to other species requires further investigation.

## Conclusions

In conclusion, our study represents a significant step forward towards the characterization of nORFs. We employed advanced statistical methods to account for low expression levels and generate a high-quality coexpression network. Despite being lowly expressed, nORFs are coexpressed with almost every cORF. We find that numerous nORFs form structured, noncanonical-only transcriptional modules which could be involved in regulating novel cellular processes. We find that many nORFs are coexpressed with genes involved in homeostasis and transport related processes, suggesting that these pathways are most likely to incorporate novel ORFs. Additionally, our investigation into the influence of genomic orientation on the expression and coexpression of *de novo* ORFs showed that ORFs located downstream of conserved ORFs are most influenced by the pre-existing regulatory environment at their locus of emergence. Our findings provide a foundation for future research to further elucidate the roles of nORFs and *de novo* ORFs in cellular processes and their broader implications in adaptation and evolution.

## Methods

### Creating ORF list

To create our initial ORF list, we utilized two sources. First, we took annotated ORFs in the *S. cerevisiae* genome R64-2-1 downloaded from SGD [103], which included 6,600 ORFs. Second, we utilized the translated ORF list from Wacholder et al. [14] reported in their *Supplementary Table 3.* We filtered to include cORFs (Verified, Uncharacterized or Transposable element genes) as well as any nORFs with evidence of translation at q value < 0.05 (Dubious, Pseudogenes and unannotated ORFs). We removed ORFs with lengths shorter than the alignment index kmer size of 25nt used for RNA-seq alignment. In situations where ORFs overlapped on the same strand with greater than 75% overlap of either ORF, we removed the shorter ORF using bedtools [104]. We removed ORFs that were exact sequence duplicates of another ORF. This left 5,878 cORFs and 18,636 nORFs, for a total of 24,514 ORFs used for RNA-seq alignment.

### RNA-seq data preprocessing

Strand specific RNA-seq samples were obtained from the Sequencing Read Archive (SRA) using the search query *(saccharomyces cerevisiae[Organism]) AND rna sequencing*. Each study was manually inspected and only studies that had an accompanying paper or detailed methods on Gene Expression Omnibus (GEO) were included. Samples were quality controlled (nucleotides with Phred score < 20 at end of reads were trimmed) and adapters were removed using TrimGalore version 0.6.4 [105]. Samples were aligned to the transcriptome GTF file containing the ORFs defined above and quantified using Salmon [106] version 0.12.0 with an index kmer size of 25. Samples with less than 1 million reads mapped or unstranded samples were removed, resulting in an expression dataset of 3,916 samples from 174 studies (Supplementary Data 1). ORFs were removed to limit sparsity and increase the number of observations in the subsequent pairwise coexpression analysis. Only ORFs that had at least 400 samples with a raw count > 5 were included for downstream coexpression analysis, n = 11,630 ORFs (5,803 canonical and 5,827 noncanonical, Supplementary Data 2).

### Coexpression calculations

The raw counts were transformed using clr. Pairwise proportionality was calculated using ρ [69] for each ORF pair. Spatial quantile normalization (SpQN) [62] of the coexpression network was performed using the mean clr expression value for each ORF as confounders to correct for mean expression bias, which resulted in similar distributions of coexpression values across varying expression levels (Supplementary Figure 2). Only ORF pairs that had at least 400 samples expressing both ORFs (at raw >5) were included. This threshold was determined empirically as detailed below.

Since zero values cannot be used with log ratio transformations, all zeros must be removed from the dataset. Proposed solutions in the literature on how to remove zeros, all of which have their pros and cons, include removing all genes that contain any zeros, imputing the zeros, or adding a pseudo count to all genes [107,108]. Removing all ORFs that contain any zeros is not possible for this analysis since the ORFs of interest are lowly and conditionally expressed. The addition of pseudocounts can be problematic when dealing with lowly expressed ORFs, for the addition of a small count is much more substantial for an ORF with a low read count compared to an ORF with a high read count [109]. For these reasons, all raw counts below 5 were set to NA prior to clr transformation. These observations were then excluded when calculating the clr transformation and in the ρ calculations. We used clr and ρ implementations in R package *Propr* [69] and implementation of SpQN from Wang et al. [62].

To determine the minimum number of samples needed expressing both ORFs in a pair, we determined the number of samples needed for coexpression values to converge within ρ ± 0.05 or ρ ± 0.1 for 2,167 nORF-cORF pairs which have a ρ > 99th percentile (before SpQN). All samples expressing both ORFs in a pair were randomly binned into groups of 10, and ρ was calculated after each addition of another sample. Fluctuations were calculated as max(ρ)-min(ρ) within a sample bin. Convergence was determined as the first sample bin with fluctuations ≤ fluctuation threshold, either 0.05 or 0.01 (Supplementary Figure 1).

### Comparing coexpression inference approaches

To compare our approach with a batch correction approach, we used clr to transform the expression matrix, followed by removing the top principal component (PC1) of the clr expression matrix to do batch correction using the function *removePrincipalComponents* from the *WGCNA* [70] R package. We then calculated ρ values and applied SpQN normalization. Additionally, we created a coexpression matrix based on TPM as well as RPKM normalized expression values instead of clr and calculated Pearson’s correlation coefficient.

### Protein Complex enrichments

We retrieved a manually curated list of 408 protein complexes in *S. cerevisiae* from the CYC2008 database by Pu et al. [64]. The coexpression matrix was filtered to contain only the 1,617 cORFs found in the CYC2008 database prior to creating the contingency table. Fisher’s exact test was used to calculate the significance of the association between coexpression and protein complex formation. Coexpressed was defined as the 99.8th ρ percentile (ρ > 0.888) considering all ORF pairs in the coexpression matrix (n = 62,204,406 ORF pairs) for Figure 1C.

### TF binding enrichments

A ChIP-exo dataset from Rossi et al. [65] containing DNA-binding information for 73 sequence-specific TFs across the whole genome was used. For each ORF we identified which TFs had binding within 200 bp upstream of the ORF’s TSS. The TSSs for all ORFs in the coexpression matrix was determined by the median 5’ transcript isoform (TIF) start positions using TIF-seq [66] dataset. Only ORFs found in the TIF-seq dataset were considered (n = 5,334 cORFs and 5,362 nORFs). To calculate the enrichments reported in Figures 1D, Supplementary Figure 5 and Supplementary Figure 7, the coexpression matrix was first filtered to only include ORFs that have at least 1 TF binding within 200 bp upstream of its TSS (n = 973 cORFs and 936 nORFs). Fisher’s exact test was used to calculate the association between coexpression and having their promoters bound by a common TF. Coexpressed was defined as the 99.8th ρ percentile (ρ > 0.888) considering all ORF pairs in the coexpression matrix (n = 62,204,406 ORF pairs) for Figure 1D.

### Coexpression matrix clustering

We used the weighted gene coexpression network analysis (*WGCNA*) package [70] in R to cluster our coexpression matrix. To do this, we first transformed our coexpression matrix into a weighted adjacency matrix by applying a soft thresholding which involved raising the coexpression matrix to the power of 12. This removed weak coexpression relationships from the matrix. We then used the topological overlap matrix (TOM) similarity to calculate the distances between each column and row of the matrix. Using the *hclust* function in R with the *ward* clustering method, we created a hierarchical clustering dendrogram. We then used the dynamic tree cutting method within the *WGCNA* package to assign ORFs to coexpression clusters, resulting in 73 clusters of which 69 were mapped to the full coexpression network. ORFs in the other four clusters were not included in the network as they did not pass the ρ threshold.

### GO analysis of clusters

We downloaded GO trees (file: go-basic.obo) and annotations (files: sgd.gaf) from ref. [110]. We used the Python package, *GOATools* [111], to calculate the number of genes associated with each GO term in a cluster and the overall population of (all) genes in the coexpression matrix. We excluded annotations based on the evidence codes ND (no biological data available). We identified GO term enrichments by calculating the likelihood of the ratio of the cORFs associated with a GO term within a cluster given the total number of cORFs associated with the same GO term in the background set of all cORFs in the coexpression matrix. We applied Fisher’s exact test and FDR with BH multiple testing correction [112] to calculate corrected p-values for the enrichment of GO term in the clusters. FDR < 0.05 was taken as a requirement for significance. We applied GO enrichment calculations only when there were at least 5 cORFs in the cluster (n=54).

### Network randomization and topology analyses

To create random networks while preserving the same degree distribution, we used an edge swapping method (Supplementary Figure 9). This involved randomly selecting two edges in the network, which were either cORF-nORF or nORF-nORF edges and swapping them. The swap was accepted only if it did not disconnect any nodes from the network and the newly generated edges were not already present in the network. We repeated this process for at least ten times the number of edges in the network. Network diameter and transitivity were calculated using R package *igraph* [113] and networks were plotted using spring embedded layout [74] in Python package *networkx* [114].

### Gene set enrichment analysis

Gene set enrichment analysis (GSEA) calculates enrichments of an ordered list of genes given a biological annotation such as GO or KEGG. For each ORF in our dataset, we used ρ values to order annotated ORFs and provided this sorted set to *fgsea* [115]. We used the GO slim file downloaded from SGD [103] for GO annotations. We used *clusterProfiler* [116] R package to download KEGG annotations using KEGG REST API [78] on 1 April 2023 and then used *fgseaMultilevel* function in *fgsea* R package to calculate enrichments for both annotations individually. To calculate GO or KEGG terms that are enriched or depleted for nORFs compared to cORFs, we calculated the number of cORFs and nORFs that had GSEA enrichments at BH adjusted FDR < 0.01. Using these counts we calculated the proportion of nORFs and cORFs associated with a GO or KEGG term and used Fisher’s exact test to assess the significance of association. P values returned by Fisher’s exact test were corrected for multiple hypothesis testing using BH correction. Odds ratios were calculated by dividing proportion of nORFs to proportion of cORFs. Proportions for the GO terms with BH adjusted FDR < 0.001 and Odds ratio greater than 2 or less than 0.5 are plotted in Figures 3A-B and are reported in Supplementary Data 5 and proportions for KEGG terms are plotted in Supplementary Figure 11 and reported in Supplementary Data 6.

### Transmembrane domain enrichment

Transmembrane domains were predicted using TMHMM 2.0 [75] for all nORFs. An ORF was classified as having a transmembrane domain if it was predicted to have at least one transmembrane domain. nORFs were classified as “coexpressed with transport-related genes” if the ORF had a GSEA enrichment at FDR < 0.01 with any of the 15 GO slim transport terms: transport, ion transport, amino acid transport, lipid transport, carbohydrate transport, regulation of transport, transmembrane transport, vacuolar transport, vesicle-mediated transport, endosomal transport, nucleobase-containing compound transport, Golgi vesicle transport, nucleocytoplasmic transport, nuclear transport, or cytoskeleton-dependent intracellular transport. Fisher’s exact test was used to calculate the significance of association between transport-related processes and transmembrane domain.

### Differential expression analysis for TF deletion and overrepresentation tests

For Hsf1 analysis, RNA-seq samples were from Ciccarelli et al. (SRA accession SRP437124) [76]. Hsf1 deletion strains were compared to wild type (WT) strains when exposed to heat shock conditions. For Sfp1 analysis, RNA-seq samples were from SRA accession SRP159150. In both cases, deletion strains were compared to WT strains. Differential expression was calculated using R package *DESeq2* [117], and ORFs were defined as differentially expressed if the log fold change (FC) in RNA expression between WT and control strains was greater than or less than 0.5 i.e. log(FC) > 0.5 or log(FC) < −0.5 and BH adjusted p-value < 0.05. ChIP-exo data for Hsf1 and Sfp1 binding was taken from Rossi et al. [65] and an ORF was labeled as having Hsf1 or Sfp1 binding if the TF was found within 200 bp upstream of the ORF’s TSS. Fisher’s exact test was performed to see if there is an association between an nORF in a GO biological process and being regulated by the TF. We define an nORF to be “in” a GO term if it has a GSEA enrichment for that GO term at FDR < 0.01. We defined an nORF as regulated by a TF if the nORF had evidence of the TF binding within 200 bp of the nORF’s TSS in ChIP-exo and has significantly downregulated expression in the TF deletion RNA-seq samples compared to the WT samples. BH p-value correction was performed for all GO terms tested. Significant GO terms and the associated regulated nORFs are reported in Supplementary Data 8.

### Detection of homologs using BLAST

We obtained the genomes of 332 budding yeasts from Shen et al. [118]. To investigate the homology of each non overlapping ORF in our dataset, we used TBLASTN and BLASTP [119] against each genome in the dataset, excluding the *Saccharomyces* genus. Default settings were used, with an e-value threshold of 0.0001. The BLASTP analysis was run against the list of protein coding genes used in Shen et al., while the TBLASTN analysis was run against each entire genome. We also applied BLASTP to annotated ORFs within the *S. cerevisiae* genome to identify homology that could be caused by whole genome duplication or transposons.

### Identification of *de* novo and conserved ORFs

To identify *de novo* ORFs, we applied several strict criteria. Firstly, we obtained translation q-values and reading frame conservation (RFC) data from Wacholder et al. [14]. All cORFs and only nORFs with a translation q-value less than 0.05 were considered as potential *de novo* candidates. We excluded ORFs that overlapped with another cORF on the same strand or had TBLASTN or BLASTP hits outside of the *Saccharomyces* genus at e-value < 0.0001. Moreover, we eliminated ORFs that had BLASTP hits to another cORF in *S. cerevisiae*. From the remaining list of candidate *de novo* ORFs, we investigated whether their ancestral sequence could be noncoding. To do this, we utilized RFC values for each species within *Saccharomyces* genus. We classified ORFs as *de novo* if the RFC values for the most distant two branches were less than 0.6, suggesting the absence of a homologous ORF in those two species. We identified conserved ORFs if a nonoverlapping cORF has an average RFC > 0.8 or has either TBLASTN or BLASTP hit at e-value < 0.0001 threshold.

To identify conserved cORFs with overlaps we first considered if the cORFs had a BLASTP outside of *Saccharomyces* genus with e-value < 0.0001. Then for two overlapping ORFs, if one had RFC > 0.8 and the other had RFC < 0.8, we considered the one with higher RFC as conserved. For the ORF pairs that were not assigned as conserved using these two criteria, we applied TBLASTN for the non-overlapping parts of the overlapping pairs. Those with a TBLASTN hit with e-value < 0.0001 were considered conserved. We found a total of 5,624 conserved ORFs and 2,756 *de novo* ORFs.

### Calculation of GO term similarities

GO term similarities were calculated using the Relevance method developed in Schlicker et al. [82]. This method considers both the information content (IC) of the GO terms that are being compared and the IC of their most informative ancestor. IC represents the frequency of a GO term; thus, an ancestral GO term has lower IC than a descendant. We used the *GOSemSim* [120] package in R that implements these similarity measures.

### Termination factor binding analysis

ChIP-exo data for Pcf11 and Nrd1 termination factor binding sites are taken from Rossi et al. [65]. This study reports binding sites at base pair resolution for *S. cerevisiae* for around 400 proteins. We used supplementary bed formatted files for Pcf11 and Nrd1, which are known transcriptional terminators, and used in house R scripts to find binding sites within the regions between the stop codon of conserved ORFs and the start codon of down same *de novo* ORFs. ORF pairs were classified as having terminators present between them if there was either Pcf11 or Nrd1 binding.

### Determining shared promoters

To determine whether two ORFs shared a promoter, we reused the TIF-Seq dataset from Pelechano et al. [66]. TIF-Seq is a sequencing method that detects the boundaries of TIFs. We extracted all reported TIFs from the supplementary data file S1 and identified all TIFs that fully cover each ORF in both YPD and galactose. We then used this information to find ORF pairs that mapped to the same TIFs for down same and up same pairs, as well as found TIFs with non-overlapping TSSs for up opposite *de novo*-conserved ORF pairs. ORF pairs where the conserved ORF was not found in the TIF-seq dataset were not included and pairs where the *de novo* ORF was not found were considered to not share a promoter.

### Web application

We utilized R language [121] and the shiny framework [73] to develop a web application which allows querying of ORFs in our dataset for information about their coexpression with other ORFs, network visualization, and GSEA enrichments. It can be accessed through a web browser and is available at https://carvunislab.csb.pitt.edu/shiny/coexpression/.

## Acknowledgments

Figures 1A, 4A, 4C, and Supplementary Figure 10 were created with BioRender.com. The authors are grateful to Dr. Aaron Wacholder, Carly Houghton, Nelson Coelho, Dr. Saurin Bipin Parikh, Jiwon Lee, Lin Chou, Alistair Turcan, Dr. Nikolaos Vakirlis, and Dr. Maria Chikina for reviewing the manuscript prior to submission.

## Author Contributions

Conceptualization: A.R., O.A., and A.-R.C.; Methodology: A.R, O.A.; Investigation: A.R, O.A.; Writing-original draft: A.R, O.A.; Writing-review and editing: A.R., O.A., and A.-R.C.; Supervision: A.-R.C. All authors approved the final version of the manuscript.

## Source code

All source codes for the analyses conducted are accessible online at https://www.github.com/oacar/noncanonical_coexpression_network

## Ethics Declarations

### Ethics approval and consent to participate

Not applicable.

### Consent for publication

Not applicable.

## Supplementary Data

Supplementary data files are available on Figshare https://doi.org/10.6084/m9.figshare.22289614

**Supplementary Data 1:** RNA-seq studies and samples used in this study. (CSV)

**Supplementary Data 2:** ORFs included in the coexpression matrix. (CSV)

**Supplementary Data 3:** Coexpression matrix generated in this study. (CSV)

**Supplementary Data 4:** GSEA analysis results for each ORF using GO BP annotations. (CSV)

**Supplementary Data 5:** List of GO BP terms that are more associated with nORFs than cORFs and statistics. (CSV)

**Supplementary Data 6:** List of KEGG terms that are more associated with nORFs than cORFs and statistics. (CSV)

**Supplementary Data 7:** GSEA analysis results for each ORF using KEGG annotations. (CSV)

**Supplementary Data 8:** GO BP terms where nORFs are regulated by either Hsf1 or Sfp1 in GO BP terms are overrepresented. (CSV)

## Supplementary Figures

**Supplementary Figure 1.**
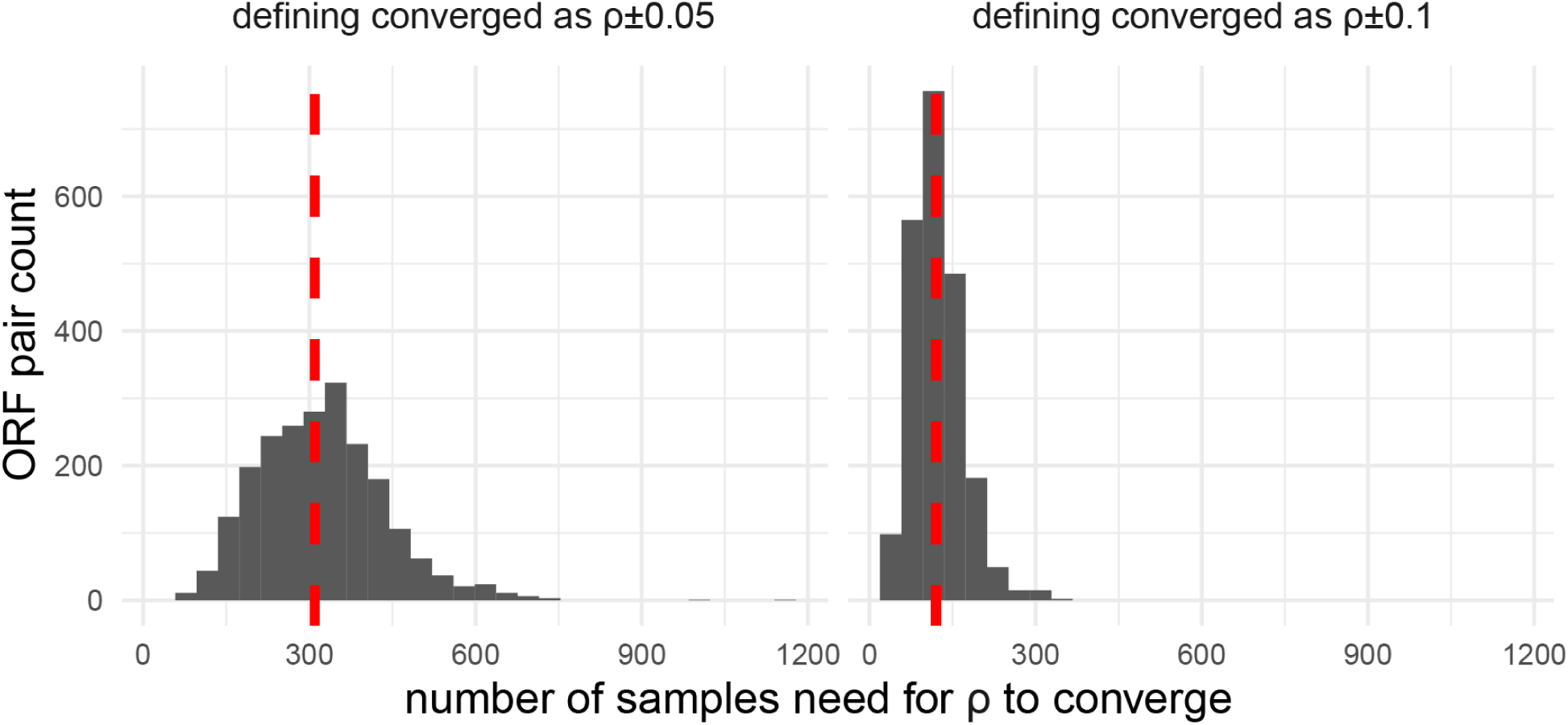
To understand the effect of sample size on coexpression values and to determine how many samples is sufficient for ρ to converge, we recalculated coexpression for a given ORF pair using n = 2 samples through n = all samples. Fluctuations were calculated as max(ρ)-min(ρ) within bins of 10 samples. The number of samples needed for ρ to converge was calculated as the first sample bin where ρ fluctuations ≤ fluctuation threshold, either 0.1 or 0.05. Histogram showing the minimum number of samples needed for ρ values to converge within ρ ± 0.05 (*left*) and ρ ± 0.1 (*right*) for 2,167 cORF-nORF pairs with ρ > 99th percentile. Red dashed lines show the median number of samples needed.

**Supplementary Figure 2.**
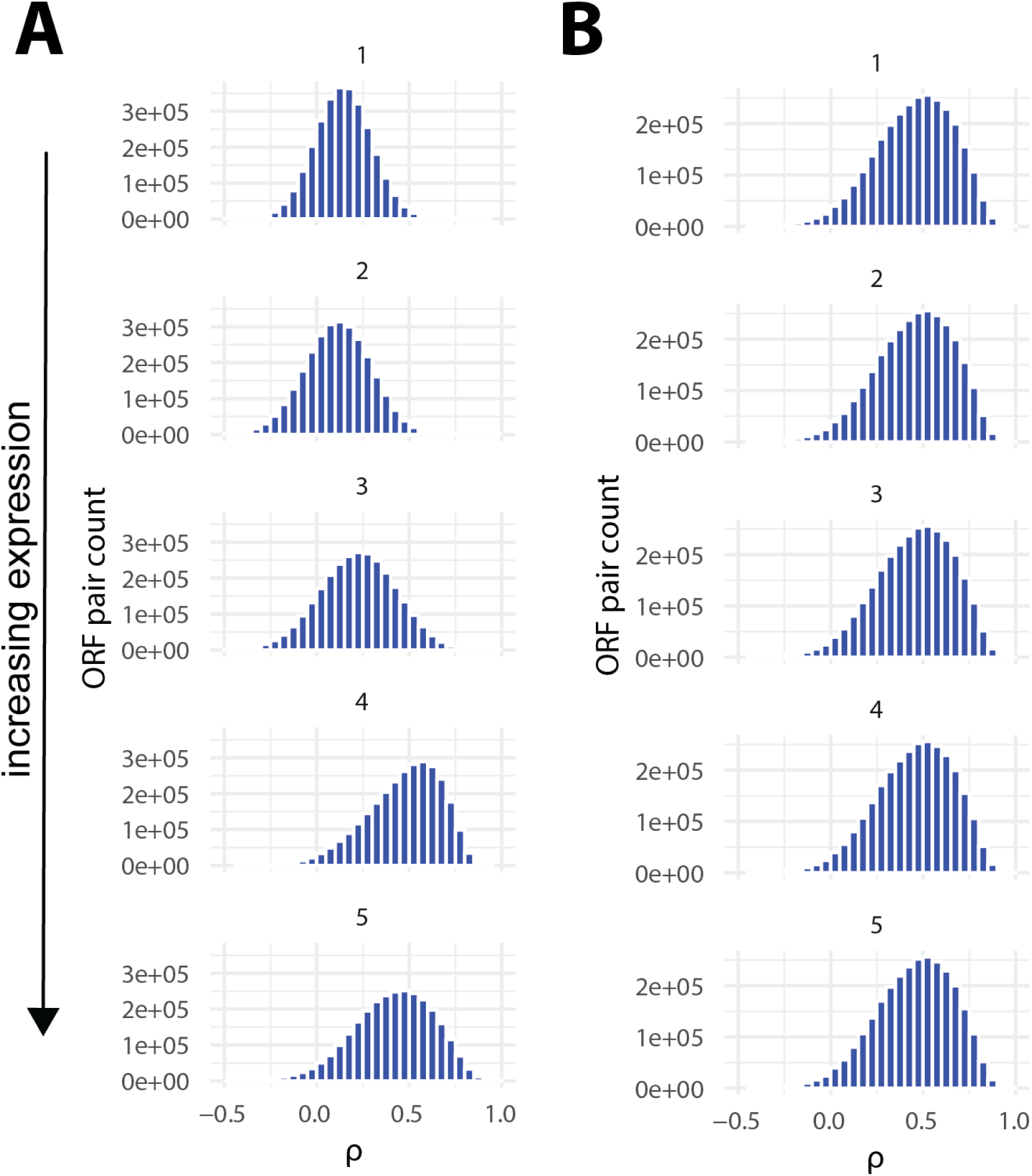
Distribution of coexpression values (ρ) for ORF pairs binned by expression level, from lowly expressed pairs *top* to highly expressed pairs *bottom*, A) before spatial quantile normalization (SpQN) and B) after SpQN, which normalizes the coexpression values so that the distribution within each expression bin is similar.

**Supplementary Figure 3.**
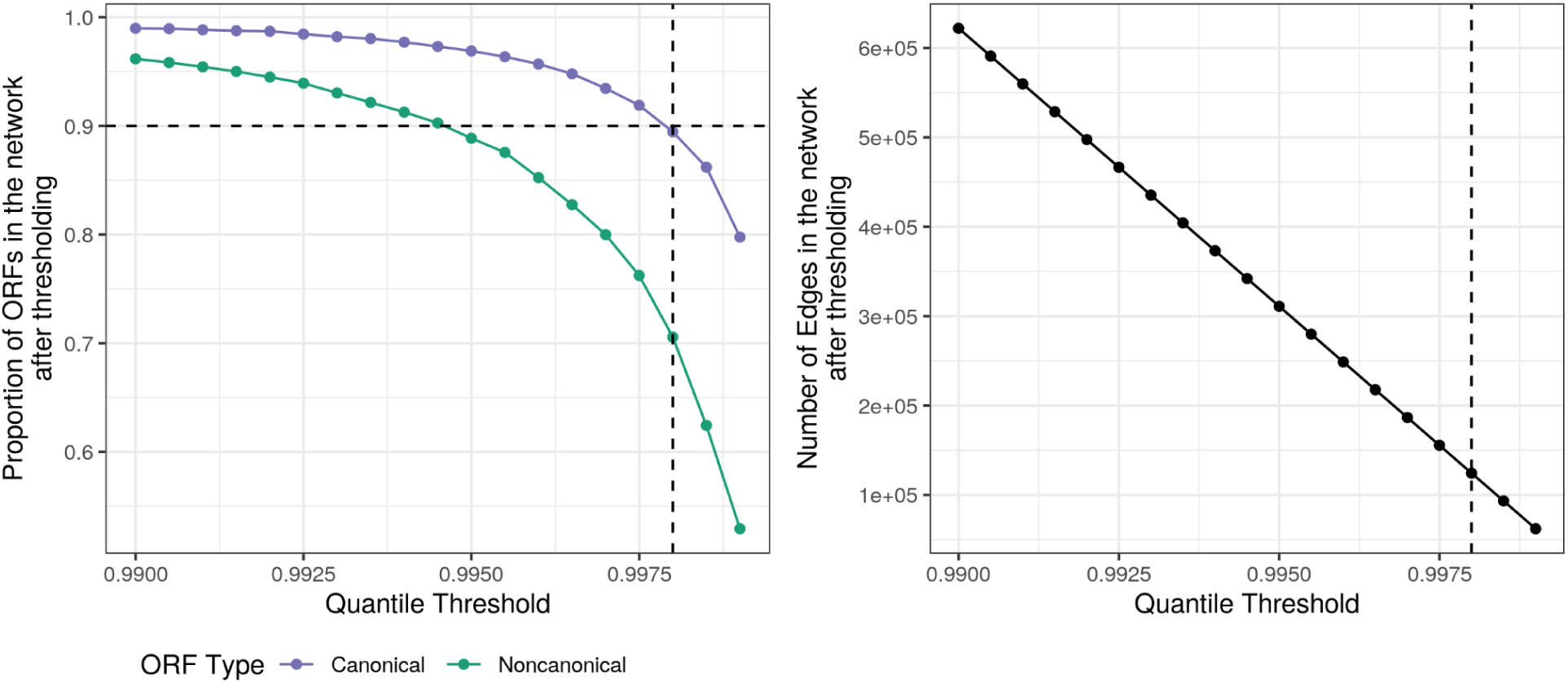
Network threshold affects cORFs and nORFs differently. *Left* shows the proportion of cORFs or nORFs in the network at each quantile threshold and the *right* shows the number of connections in the network. Dashed line represents 0.9998 quantile which was chosen for creating the network.

**Supplementary Figure 4.**
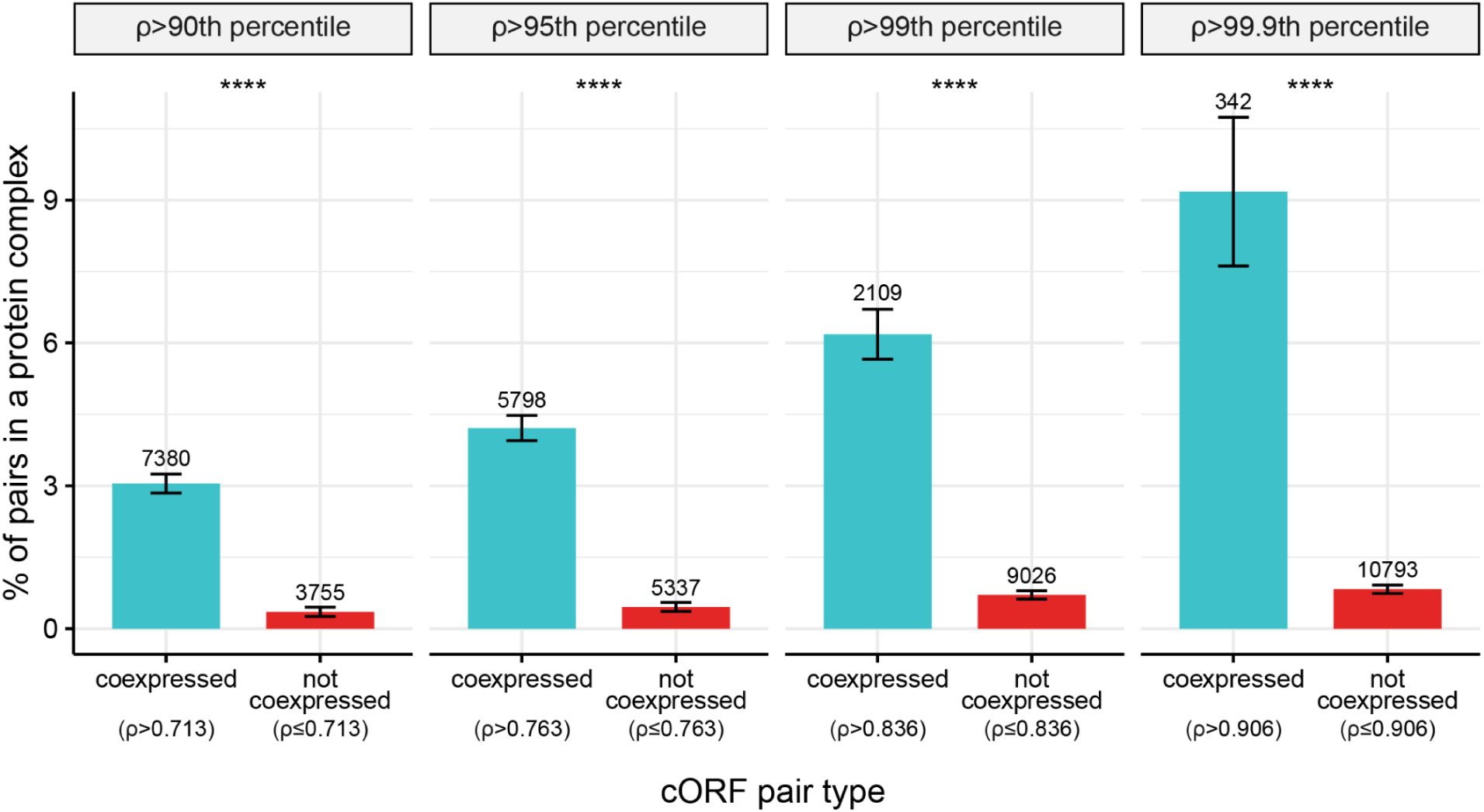
Coexpressed cORFs pairs are more likely to encode proteins that form protein complexes than non-coexpressed cORF pairs, and this is consistent across different coexpression cutoffs. Coexpression was defined using the top 90th, 95th, 99th, and 99.9th percentile of all ORF pairs in the network (n = 62,204,406 ORF pairs). 90th percentile (ρ > 0.713) Odds ratio = 8.89; 95th percentile (ρ > 0.763) Odds ratio = 9.59; 99th percentile (ρ > 0.836) Odds ratio = 9.23; 99.9th percentile (ρ > 0.906) Odds ratio = 12.1; Fisher’s exact test p < 2.2e-16 for all comparisons. Numbers above bars represent the number of ORF pairs in each category. Error bars represent the standard error of the proportion. A list of 408 protein complexes were retrieved from Pu et al. CYC2008 database [64]. Enrichments were calculated using only the 1,617 cORFs found in the CYC2008 database.

**Supplementary Figure 5.**
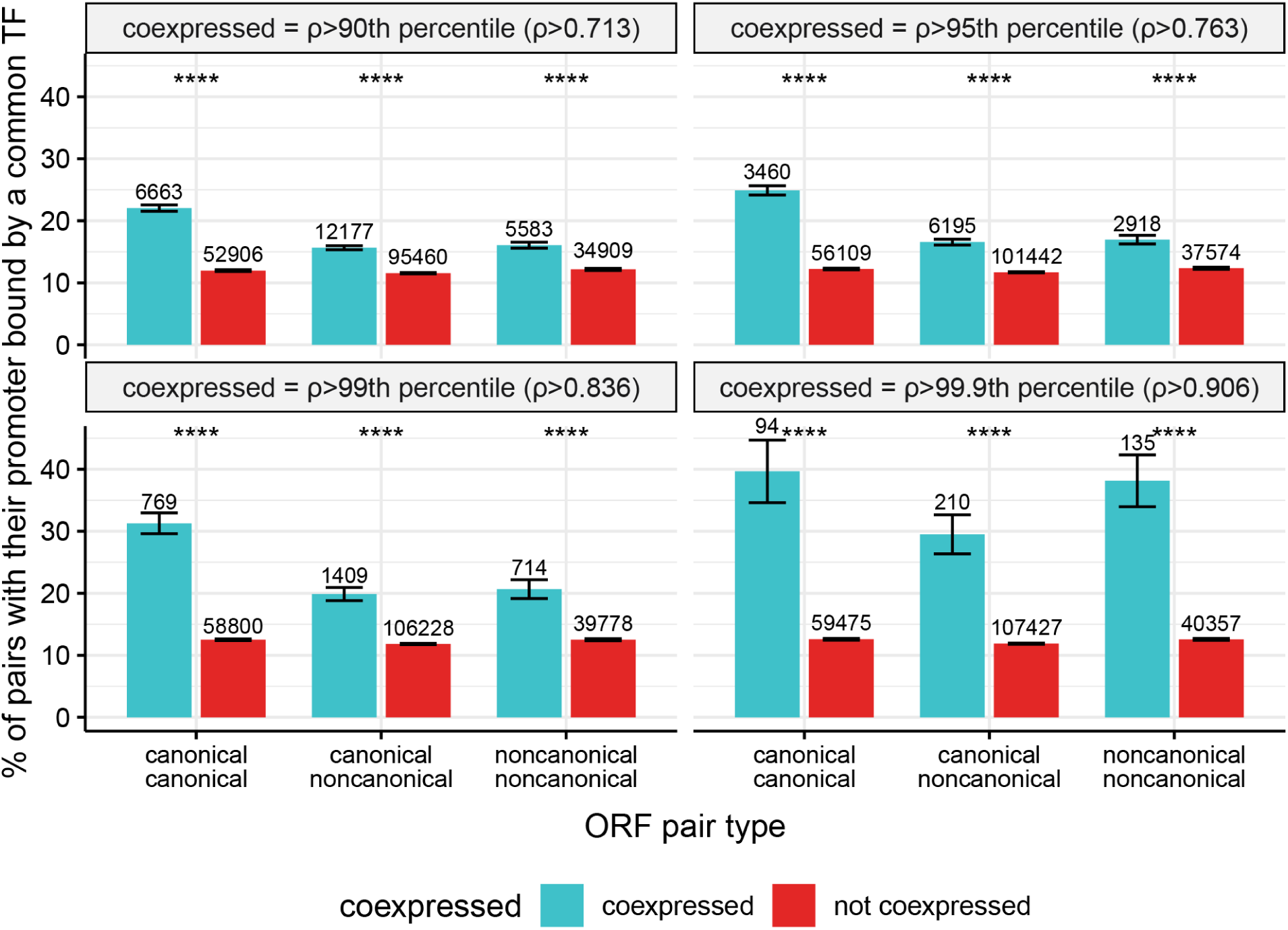
Coexpressed ORF pairs are more likely to have their promoters bound by a common TF than non-coexpressed ORF pairs, and this is true across different coexpression cutoffs and for canonical-canonical (cc), canonical-noncanonical (cn) and noncanonical-noncanonical (nn) ORF pairs. Coexpression was defined using the top 90th, 95th, 99th, and 99.9th percentile of all ORF pairs in the network (n = 62,204,406 ORF pairs). 90th percentile (ρ > 0.713): cc Odds ratio = 2.08, cn Odds ratio = 1.42, nn Odds ratio = 1.38; 95th percentile (ρ > 0.763): cc Odds ratio = 2.38, cn Odds ratio = 1.50, nn Odds ratio = 1.45; 99th percentile (ρ > 0.836): cc Odds ratio = 3.19, cn Odds ratio = 1.85, nn Odds ratio = 1.82; 99.9th percentile (ρ > 0.906): cc Odds ratio = 4.57, cn Odds ratio = 3.10, nn Odds ratio = 4.29; ****: Fisher’s exact test p < 2.2e-16 for all comparisons. Error bars represent the standard error of the proportion. Using a ChIP-exo dataset from Rossi et al. [65] containing DNA-binding information for 73 sequence-specific TFs, TF binding was defined as a ChIP-exo peak within 200 bp upstream of the ORF’s TSS. Only ORFs whose promoter was bound by at least one TF were considered. Numbers above bars represent the number of ORF pairs in each category.

**Supplementary Figure 6.**
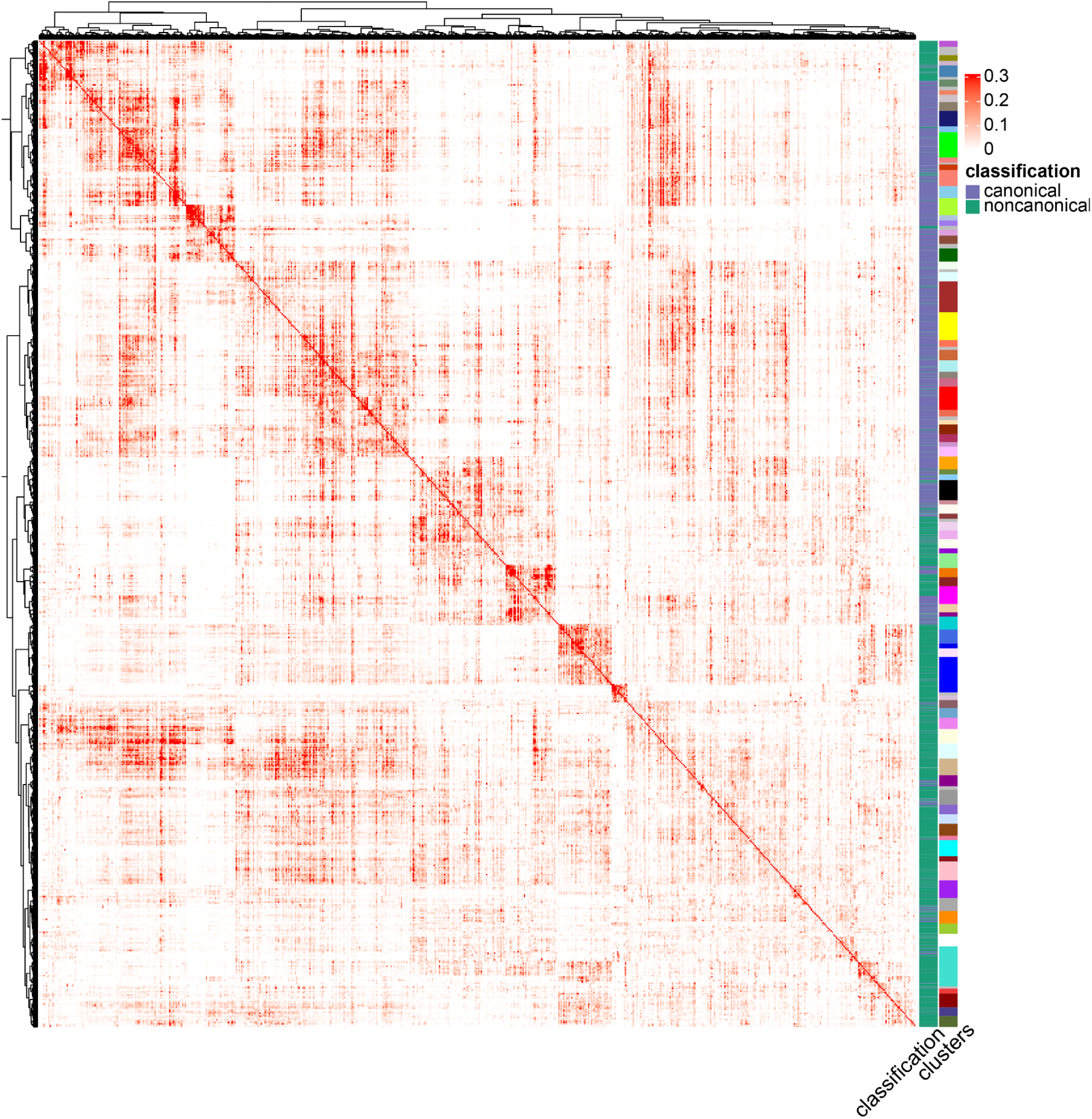
Clustered matrix heatmap. Coexpression values are first transformed by taking power of 12 and then WGCNA pipeline [70] is applied. Clusters are determined by cutting dendrograms (see methods for details). Colors on ‘clusters’ section represent the different clusters. Values of 0.3 and above are represented by red to show the structure of the heatmap.

**Supplementary Figure 7.**
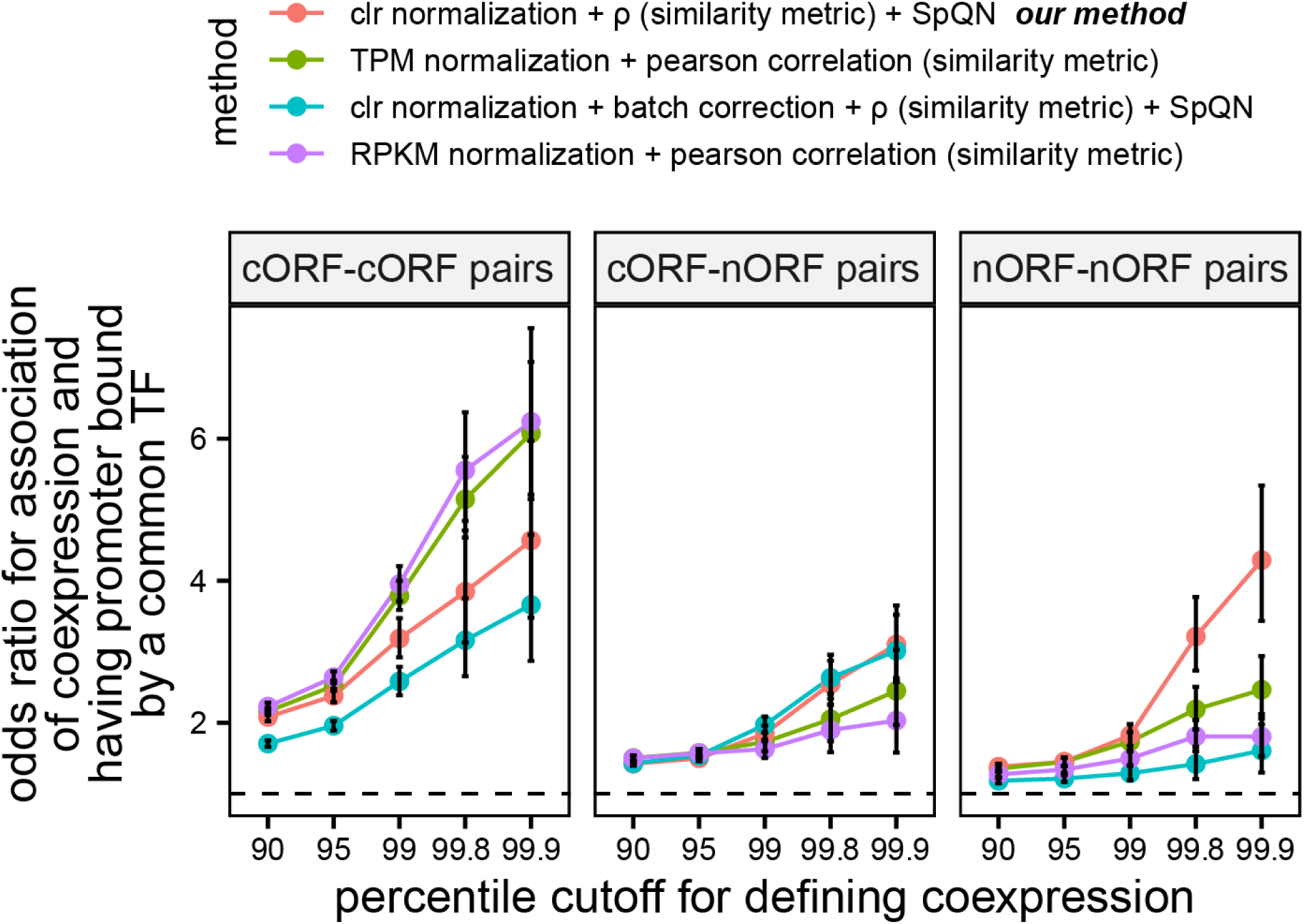
Using clr normalization, ρ similarity metric and SpQN normalization leads to the highest odds ratios for nORF-nORF coexpressed pairs to also have their promoters bound by common TFs. Our method (*pink*) uses clr to transform the expression matrix, uses proportionality metric ρ to calculate coexpression and SpQN to normalize the coexpression matrix. Method TPM + pearson (*green*) uses TPM to normalize the expression matrix followed by Pearson correlation to calculate coexpression. Method clr + batch correction + rho + SpQN (*blue*) uses clr to transform the expression matrix, followed by removing the top principal component of the clr expression matrix to do batch correction, followed by calculating coexpression using proportionality metric ρ and SpQN normalization of the coexpression matrix. Method RPKM + pearson correlation (*purple*) uses RPKM to normalize the expression matrix followed by Pearson correlation to calculate coexpression. Coexpression percentiles were determined using all ORF pairs (n = 62,204,406 ORF pairs). All odds ratios are significant at p < 2.15e-5, Fisher exact test. Batch correction performed by removing the top principal component on the clr transformed expression matrix. Error bars represent the 95% confidence interval of the odds ratio. Dashed line shows an odds ratio of 1.

**Supplementary Figure 8.**
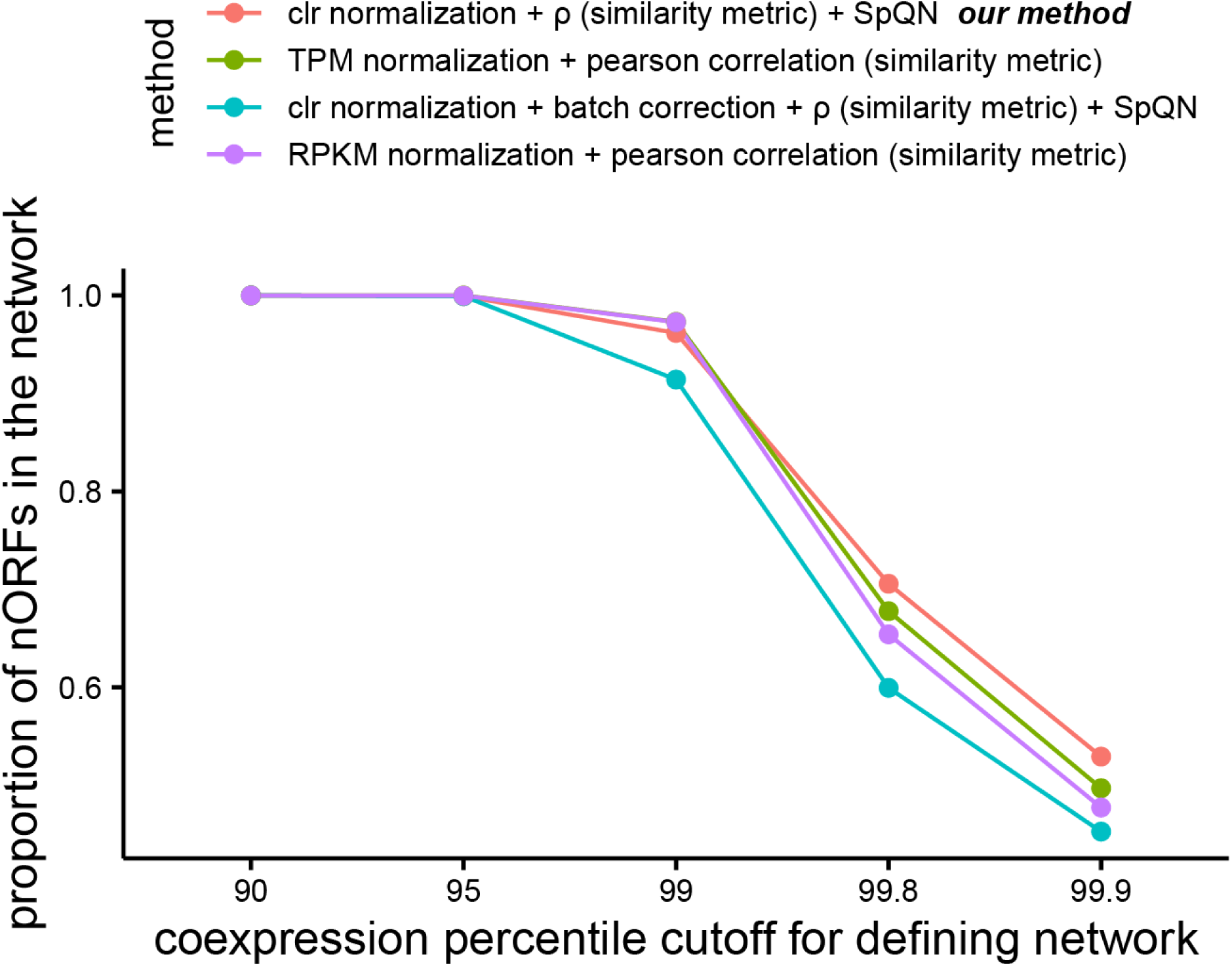
Proportion of nORFs defined as coexpressed (and therefore included in the coexpression network) at various coexpression percentile cutoffs using four different methods. Our method (*pink*) uses clr to transform the expression matrix, uses proportionality metric ρ to calculate coexpression and SpQN to normalize the coexpression matrix. Method TPM + Pearson (*green*) uses TPM to normalize the expression matrix followed by Pearson correlation to calculate coexpression. Method ρ + batch correction (*blue*) uses clr to transform the expression matrix, followed by removing the top principal component of the clr expression matrix to do batch correction, followed by calculating coexpression using proportionality metric ρ and SpQN normalization of the coexpression matrix. Method RPKM + pearson correlation (*purple*) uses RPKM to normalize the expression matrix followed by Pearson correlation to calculate coexpression. Coexpression percentiles were determined using all ORF pairs (n = 62,204,406 ORF pairs).

**Supplementary Figure 9.**
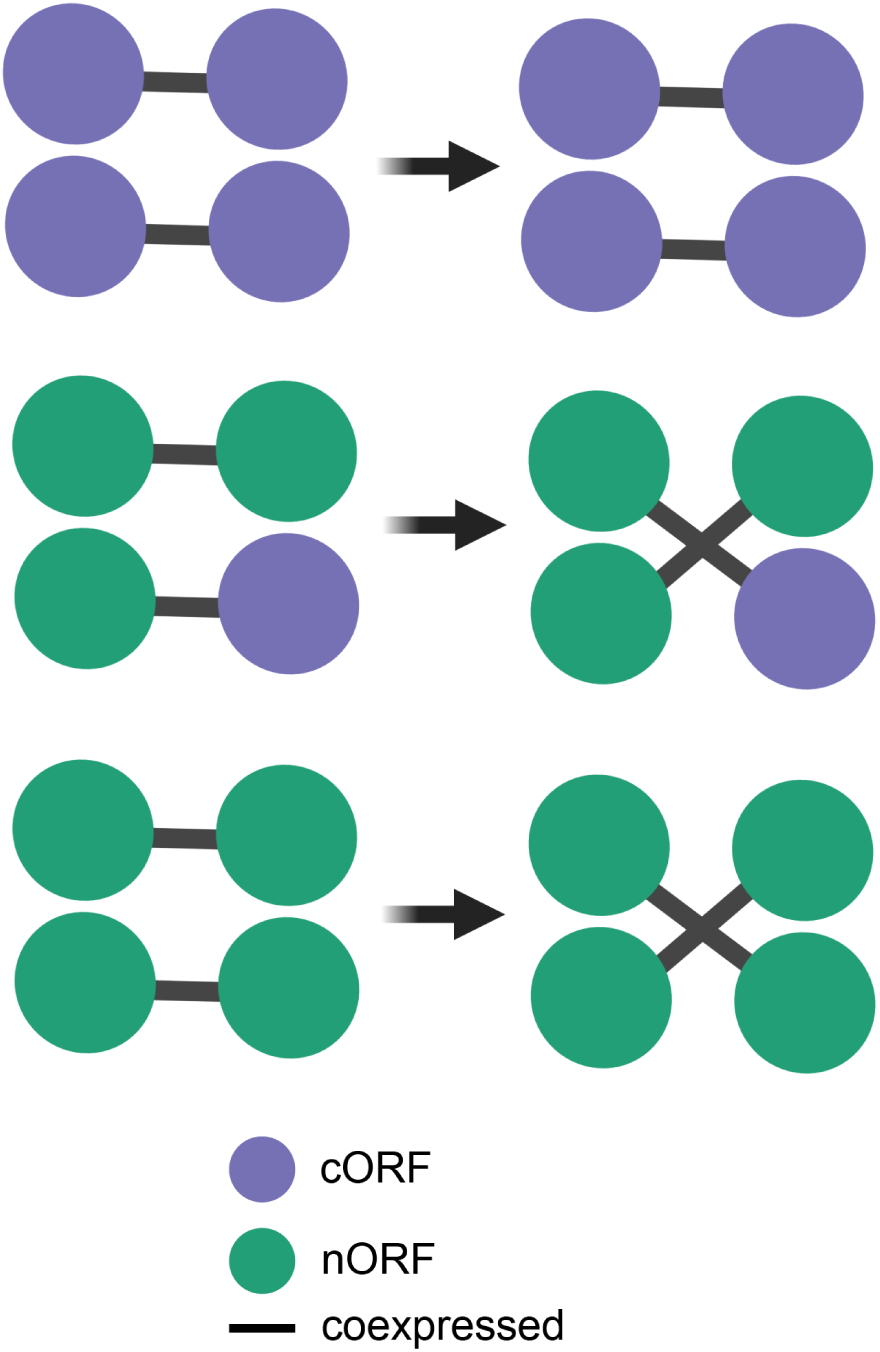
Strategy for generating randomized networks. Edges between cORF-nORF and nORF-nORF pairs were swapped in a pairwise manner such that the degree of each node stayed the same. Edges between cORF-cORF pairs were not randomized.

**Supplementary Figure 10.**
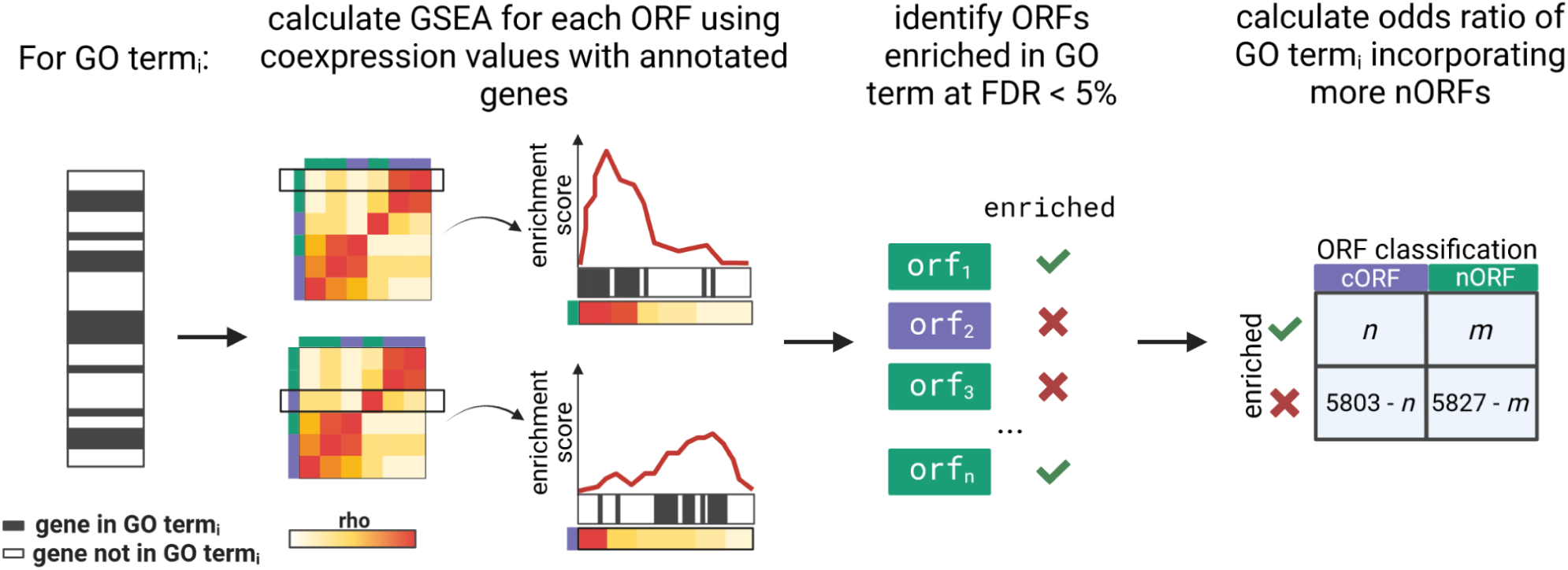
GSEA pipeline using coexpression profiles to find GO terms that are more likely to incorporate nORFs.

**Supplementary Figure 11.**
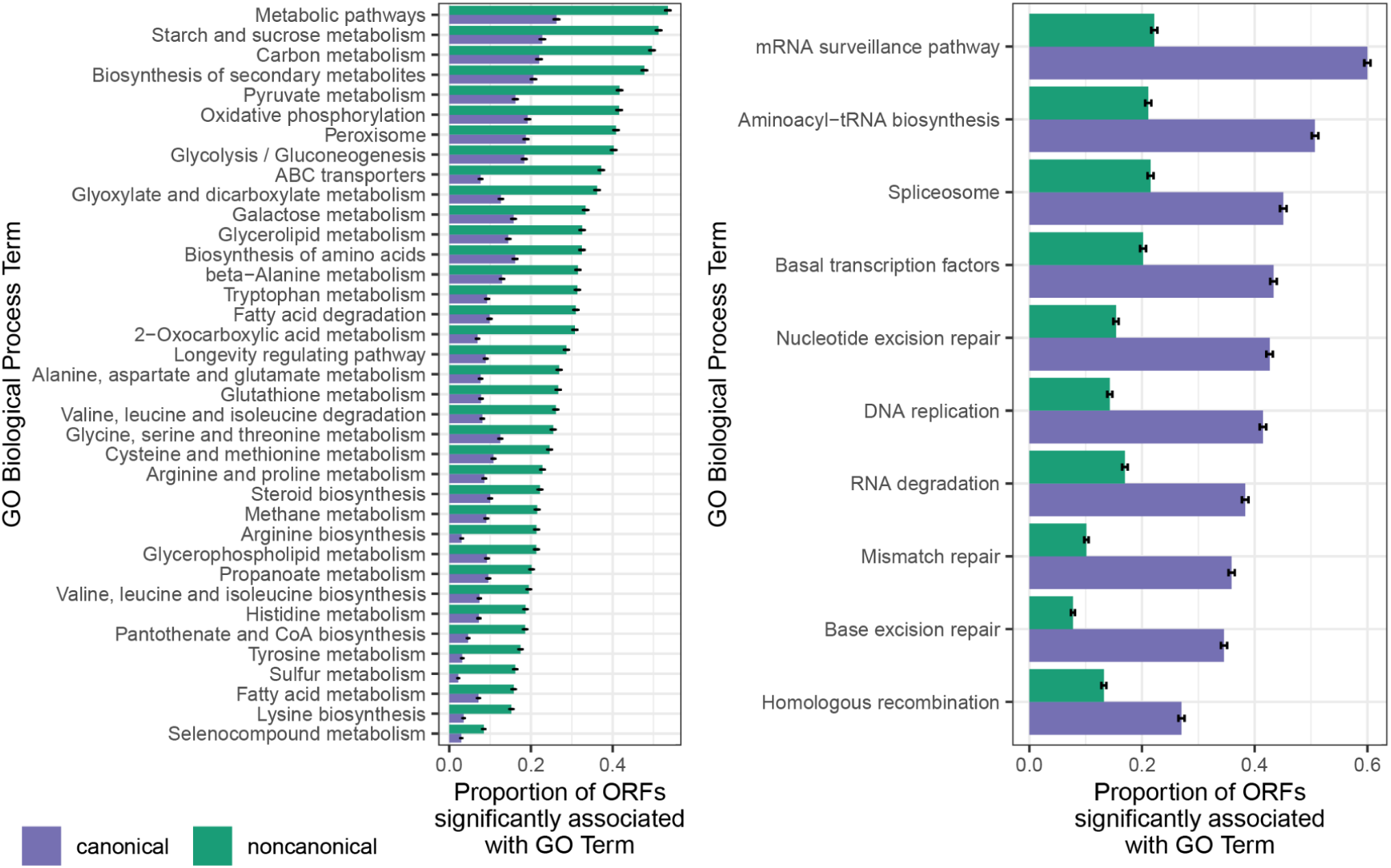
KEGG pathways that proportionally have more (*left*) (Odds ratio > 2, n = 37 terms) or less (*right*) (Odds ratio < 0.5, n = 10 terms) GSEA enrichments with nORFs compared to cORFs (y-axis ordered by nORF enrichment proportion from highest to lowest, BH adjusted FDR < 0.001 for all terms, Fisher’s exact test). Error bars represent the standard error of the proportion.

**Supplementary Figure 12.**
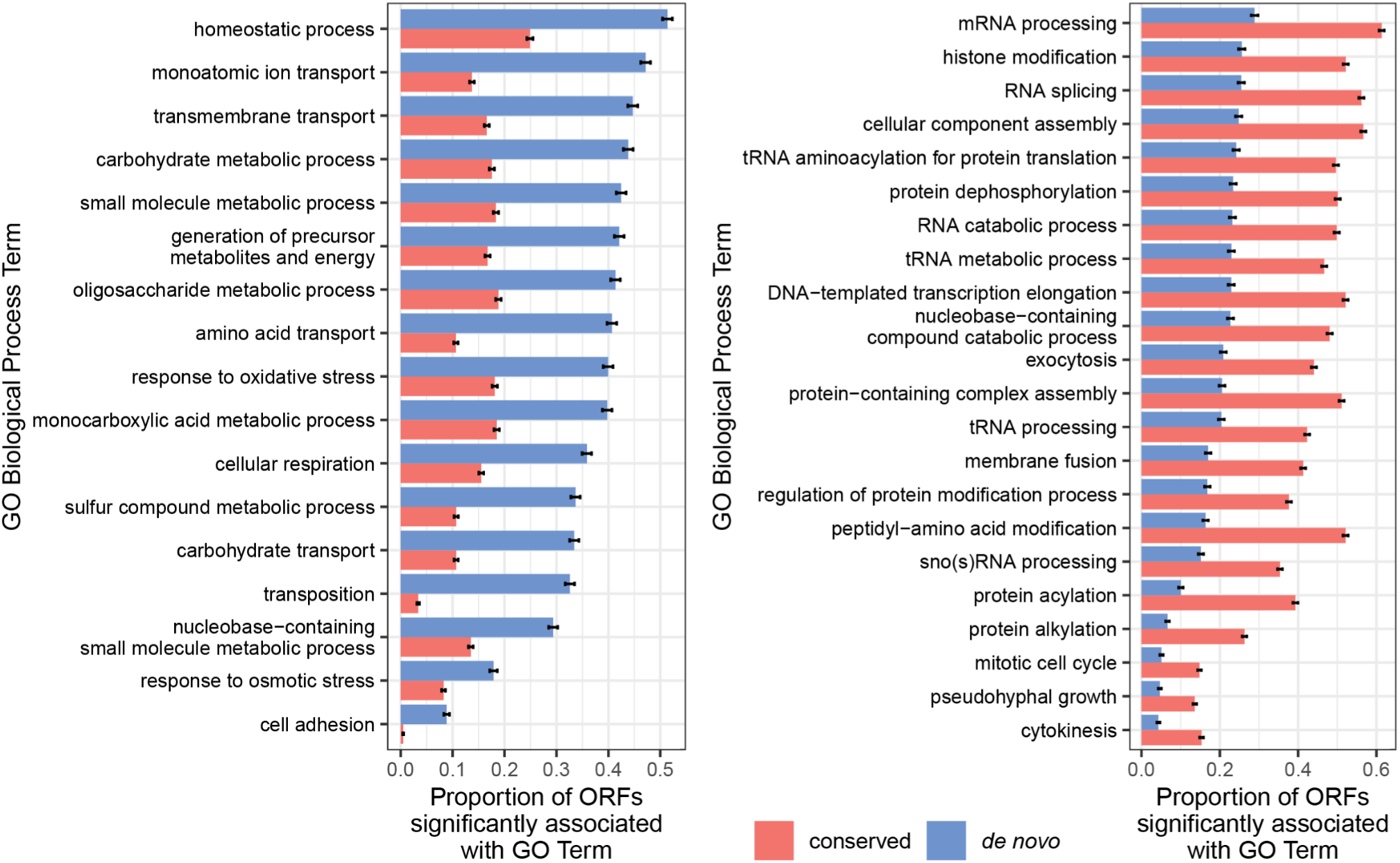
GO terms that proportionally have more (*left*) (Odds ratio > 2, n = 35 terms) or less (*right*) (Odds ratio < 0.5, n = 11 terms) GSEA enrichments with *de novo* ORFs compared to conserved ORFs (y-axis ordered by *de novo* ORF enrichment proportion from highest to lowest, BH adjusted FDR < 0.001 for all terms, Fisher’s exact test). Error bars represent the standard error of the proportion.

**Supplementary Figure 13.**
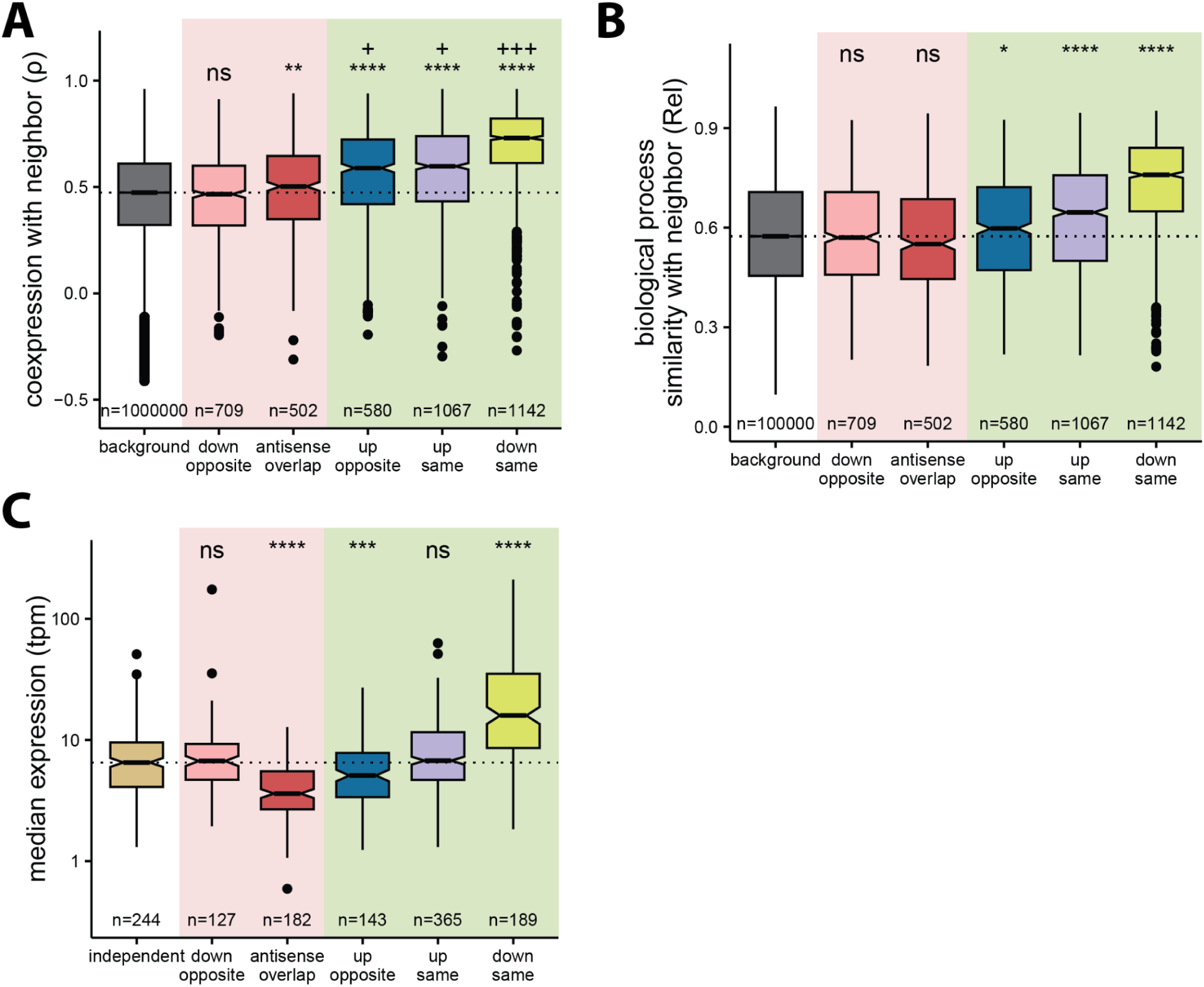
A) Coexpression (y-axis) of *de novo* ORFs with neighboring conserved ORFs per orientation (x-axis). Down same *de novo* ORFs tend to be highly coexpressed with their neighbors; background: *de novo*-conserved ORF pairs located on different chromosomes. B) Biological process similarity (y-axis) of *de novo* ORFs with neighboring conserved ORFs per orientation (x-axis). Similarity measured by calculating semantic similarity between GSEA enrichments for neighboring *de novo*-conserved ORF pairs using relevance metric (0 = no similarity, 1 = perfect overlap); background: *de novo*-conserved ORF pairs located on different chromosomes. C) Median expression of *de novo* ORFs (y-axis) per orientation (x-axis). *De novo* ORFs located downstream on the same strand as conserved ORFs have the highest expression among different orientations (considering only ORFs in only a single orientation, dashed box in panel 4D; independent: *de novo* ORFs located further than 500 bp from all conserved ORFs). For panels A-B-C: Mann-Whitney U-test, ****: p ≤ 0.0001, ***: p ≤ 0.001, **: p ≤ 0.01, *: p ≤ 0.05, ns: not-significant, +: small effect size (Cliff’s d < 0.33), ++: medium effect size (Cliff’s d < 0.474), +++: large effect size (Cliff’s d ≥ 0.474); all orientations are compared to either background pairs (A, B) or to independent ORFs (C).

**Supplementary Figure 14.**
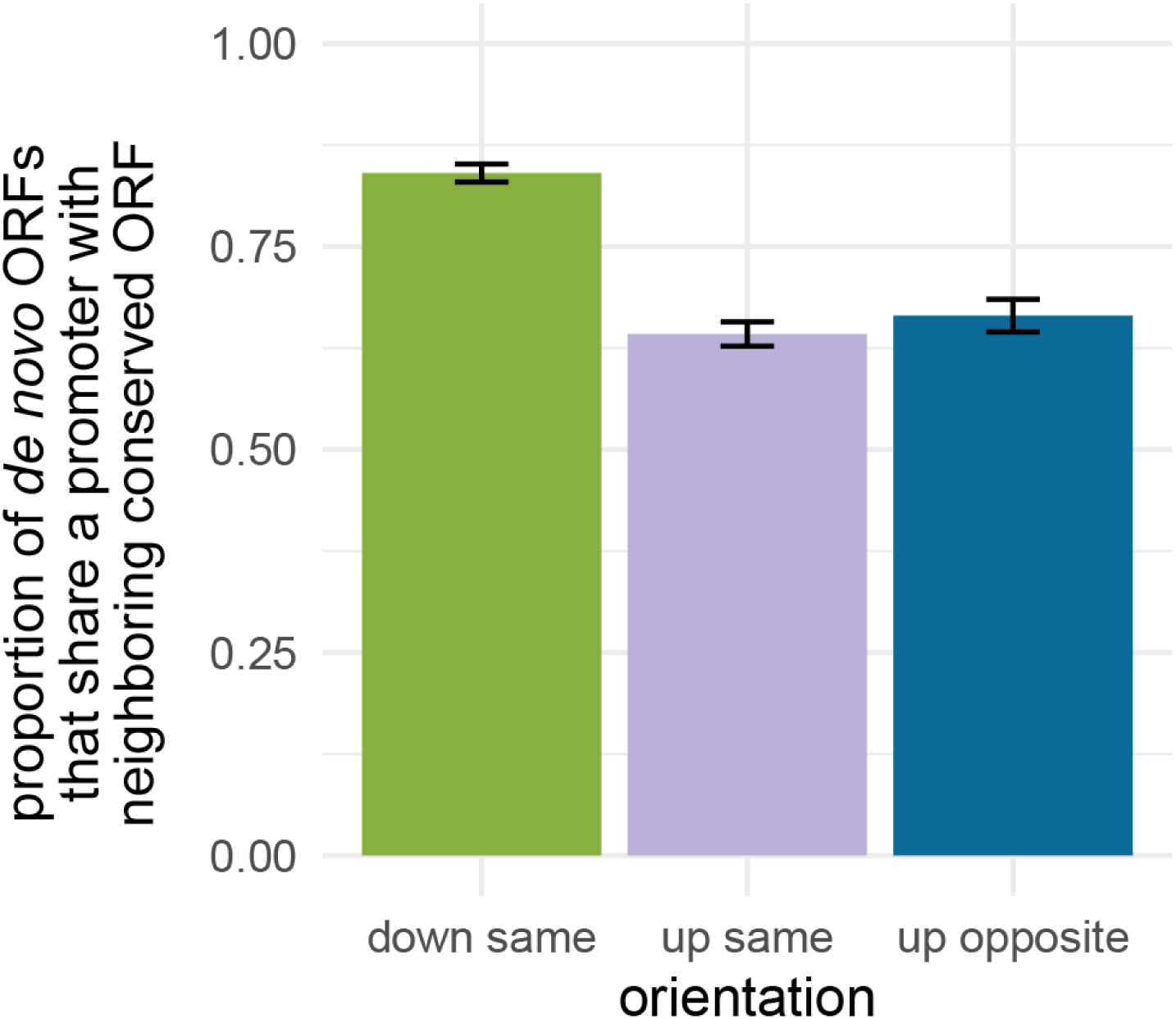
Proportion of *de novo* ORFs that share a promoter with their neighboring conserved ORF. To determine if ORFs shared a promoter with neighbors we used a publicly available TIF-seq dataset from Pelechano et al [65]. We defined down same or up same ORFs as sharing a promoter if they mapped to the same transcript at least once, and defined up opposite ORFs as sharing a promoter if their respective transcripts did not have overlapping TSSs. We found that 84% of down same (n = 174), 64% of up same (n = 368), and 66% of up opposite (n = 185) *de novo* ORFs share a promoter with their neighboring conserved ORF. Error bars represent the standard error of the proportion.

